# Extra-ciliary role for polycystins in regulation of Ezrin and renal tubular morphology

**DOI:** 10.1101/2025.11.05.684799

**Authors:** Eryn E. Dixon, Ava M. Zapf, Victoria L. Halperin Kuhns, Denis Basquin, Rachel M. Park, Richard Coleman, Lidiya Franklin, Allison C. Lane-Harris, Alexis Hofherr, Michael Köttgen, Feng Qian, Paul A. Welling, Terry J. Watnick, Owen M. Woodward

## Abstract

Full understanding of the functions of the polycystin proteins, PC1 and PC2, in renal epithelial cells is obscured by signaling complexity and renal injury that occurs in Autosomal Dominant Polycystic Kidney Disease (ADPKD). The polycystins likely function as a complex in the primary cilium, yet previous work hinted at a critical role for PC1 function outside of the primary cilium (extra-ciliary) during tubule development. Here, we investigate an extra-ciliary role for the polycystins in regulating renal cell and tubular morphology. First, we found acute loss of polycystins significantly increased the circularity of renal epithelial cells and tubuloids grown in 3D culture. Next, we demonstrated that both PC1 and PC2 can immunoprecipitate Ezrin, an ERM protein important for apical compartment shape. In human ADPKD renal cystic tissue, and after acute inducible knockout of *Pkd1* or *Pkd2,* we found that Ezrin protein abundance is significantly reduced, with the remaining Ezrin protein mis-localized. Immunofluorescence in 2D cells and 3D tubuloids suggested acute polycystin loss specifically reduced the active form of Ezrin at the apical surface, leaving inactive Ezrin colocalized with ZO1 in the cell junctions. A specific ERM phosphorylation inhibitor, NSC668394, phenocopied the increased circularity observed in the *Pkd1* knockout spheroids, as did inhibition of PKC activity, implicating the polycystin complex in regulating Ezrin phosphorylation. Our data strongly support a role for the polycystin complex in regulating renal cell and tubular shape via interactions with the ERM protein Ezrin, interactions that do not require trafficking to the primary cilium.

## Introduction

Autosomal dominant polycystic kidney disease (ADPKD) is the most common inherited renal disease and a major contributor to the population burden of end stage kidney disease (ESKD)^1–3^. ADPKD is caused by mutations in either *PKD1* or *PKD2*. The *PKD* gene products, polycystin-1 (PC1) and polycystin-2 (PC2), are hypothesized to form a signaling complex at the membrane, localizing to the primary cilium, and possibly cell-cell junctions, and endoplasmic reticulum (ER)^4^. PC1 serves as the receptor with reported roles in ligand or flow detection in the cilium^1,5^. PC2 is a TRP-like nonselective cation channel that transduces signals possibly using calcium as its messenger^6^. Despite decades of investigation, we still do not fully understand the physiological function of the PC1/PC2 complex.

Strong evidence supports a critical role for polycystin complex function in the primary cilia^7–9^, but there also exists growing evidence for an extra-ciliary role for the polycystins^10–13^. Tissue and subcellular localization of the polycystins, especially PC1, is complicated by very low expression levels in the adult kidney, challenging 3D folded structure, and a complex pattern of cleavage making antibody based subcellular localization outside of the cilium challenging^1^. In contrast, experiments focused on describing PC1/PC2 complex function using gene oblation / knockout have identified several extra-ciliary functions mediated by the polycystin proteins^1,11–14^.

Trafficking of PC1 and PC2 to the primary cilium requires both physical interaction between PC1 and PC2 and auto-proteolytic cleavage of PC1 at the G protein-coupled receptor proteolytic cleavage site (GPS) at the base of the extracellular domain^5,15,16^. Investigations of endogenous PC1 protein demonstrate that although most of the PC1 molecule exists as the cleaved version in the cilium, there is a significant population of un-cleaved PC1, excluded from ciliary trafficking ^17^. Exactly what the role of GPS cleavage and extra-ciliary PC1 function is remains a critical gap in our understanding of polycystin function^1,15,16,18,19^. While germline knockout of *Pkd1* in mouse models produces a lethal embryonic phenotype of massively cystic kidneys, with severe vascular, CNS, bone, and lung phenotypes^20^, mutations in the GPS site of mouse PC1 (T3041V)^19^ that prevent cleavage and ciliary localization, result in a normal phenotype at birth^18^. Though these *Pkd1*V/V mice subsequently develop renal cysts, the other tissue phenotypes fail to emerge^21^, supporting a functional role for the un-cleaved, extra-ciliary PC1^18^ conserved across tissue types.

Unbiased approaches to better understand the proteins that interact with PC1 have found most of the PC1 “interactome” proteins are non-ciliary^12,22^. Tagged PC1 in a cell model used as bait identified key actomyosin regulatory proteins as PC1 interactors, demonstrating a role for PC1 in regulating actomyosin contraction, cell stiffness, and YAP activation^12^. Additionally, inducible knockout of *Pkd2* in mouse renal epithelial cells demonstrated the loss of PC2 results in altered expression of many genes related to cell junctions, matrix interactions, actin cytoskeleton, and cell morphology^14^ including TNS1 (Tensin1), which encodes an important linker of actin to beta-integrins on the basolateral membrane^14^. A more recent effort has described the *in vivo* mouse brain PC1 interactome reporting a total of 817 interactors of which 27 were ciliary proteins, whereas 200 were cytoskeletal and junctional proteins^22^, data which further supports a non-ciliary function of PC1/PC2 conserved across different tissue types.

Finally, understanding where and how the polycystins function is ultimately dependent on our understanding of their contribution to cystogenesis. The mechanism of cystogenesis associated with ADPKD can be conceptually parsed into several key components, including loss of polycystin protein, changes in cell and tubule shape, increases in proliferation, and fluid secretion/accumulation^14,23^. In practice, experimentally teasing apart the stages of cystogenesis is challenging, but necessary to understanding where and when the polycystins function is required. In this study, to better understand how extra-ciliary polycystin signaling complex regulates the processes of cystogenesis, we have used conditional inactivation of *Pkd1/Pkd2* in a 3D-tubuloid model to measure acute changes in cell and tubuloid shape after *Pkd1/2* knock out. We found that acute loss of polycystins alters cell and tubular morphology, and these alterations precede increases in tubuloid size. We show PC1/PC2 interact with the cytosolic ERM protein Ezrin, and mechanistically this interaction is crucial for the maintenance of tubuloid shape. Further, we demonstrate PC1/PC2 regulates the activation of Ezrin and its subsequent localization to the apical membrane in renal epithelial cells, potentially by facilitating interactions with PKC isoforms. These findings strongly support a role for the polycystin complex in regulating renal cell and tubular shape via interactions with the ERM protein Ezrin, interaction that do not require trafficking to the primary cilium, and do not alter cell cycle or proliferation.

## Results

### Polycystin expression regulates cell and tubular morphology in 3D cultured renal epithelial cells

The slowly progressive nature of ADPKD has made understanding the most proximate physiological changes after the loss of polycystin function extremely difficult to interpret. To better understand the earliest alterations in cellular function and morphology after polycystin loss, we created unique immortalized mouse renal epithelial cell lines that allow for inducible *Pkd1* and *Pkd2* knockout with isogenic controls (Supplementary Figure 1). We cultured immortalized mouse *Pkd1^fl/fl^ Pax8rtTA, TetO-Cre* medullary (M) renal epithelial cells (*Pkd1(M)* = clone #313); after DMSO (*Pkd1+(M))* or doxycycline (*Pkd1-(M))* treatment in Matrigel using a “sandwich” technique with the GDNF pulse^14^ that allows cell to form into tube-like shapes, or tubuloids, within 14 days. We quantified two structural characteristics: circularity, to capture cell and tubular morphology; and tubuloid size (area) as a proxy for rate of structural growth and cell proliferation (using a semiautomated quantification protocol and FIJI). After 14 days, the DMSO treated tubuloids resembled complicated, branching, truncated tubules (Figure 1A). In contrast, the doxycycline treated cells grow in to large, transparent, spherical structures, consistent with the loss of polycystin function. The *Pkd1-(M)* structures were significantly larger (Figure 1B, p<0.05), and significantly more circular than the *Pkd1+(M)* tubuloids (Figure 1C, p<0.05). These 14 day phenotypes are consistent with previous work showing that polycystin loss leads to eventual large cysts with altered morphology and increased cell proliferation^14^. To investigate if these two pathways could be decoupled we looked at earlier time points to understand if changes in size and shape occur simultaneously or consecutively. After 6 days in 3D culture (Figure 1D), both the DMSO and doxycycline treated cultures produced numerous small tubuloids. We again quantified tubuloid size (area) and circularity and found that at the earlier time point the respective size of the structures was not significantly different (Figure 1E), yet the *Pkd1-(M)* tubuloids were already significantly more circular (Figure 1F, p<0.05) than the control tubuloids. We then sought to understand if there was a relationship between size and circularity using a linear regression analysis (Figure 1G). In the control *Pkd1+ (M)* tubuloids there is a tight negative correlation between circularity and size: the larger the structure, the less circular they became. In contrast, the *Pkd1-(M)* tubuloids are more circular regardless of size, and the correlation is significantly weaker than the controls (significant difference in regression slope, p<0.0001). To determine if we could observe these differences in the relationship between size and circularity in real time we repeated our 14 day 3D culture experiment with the same *Pkd1+/−(M)* tubuloids, and tracked individual structures (photographed every day) from day 7 until the end of the experiment on day 14 (Figure 1H,I). Mirroring what we observed at the population level, individual control tubuloids became less circular during the 7 day tracking period (Figure 1J), whereas the *Pkd1-(M)* structures became significantly more circular (p<0.05). One major attribute that contributed to increased circularity was the significant decrease in branches of the *Pkd1-(M)* tubuloid/ spheroids at day 14 (Figure 1K, p<0.05). Next, to better understand the mechanisms underlying the changes in tubuloid characteristics after *Pkd1* loss, we cultured *Pkd1 (M)* (clone #313) cells after DMSO (*Pkd1+(M))* or doxycycline (*Pkd1-(M))* treatment in Matrigel for 14 days, stained for ZO1 (Figure 1L), and then measured the individual cell shape in the tubloids (Figure 1M). We found the *Pkd1-* tubuloids contained cells with a significantly smaller aspect ratio (the major and minor axis are closer in size, p<0.05), reflective of a more circular cell shape. These studies strongly support a role for the polycystins in the 1^st^ order control of cell and tubular morphology, preceding any 2^nd^ order alterations in cell proliferation or altered cycle.

**Figure 1:**
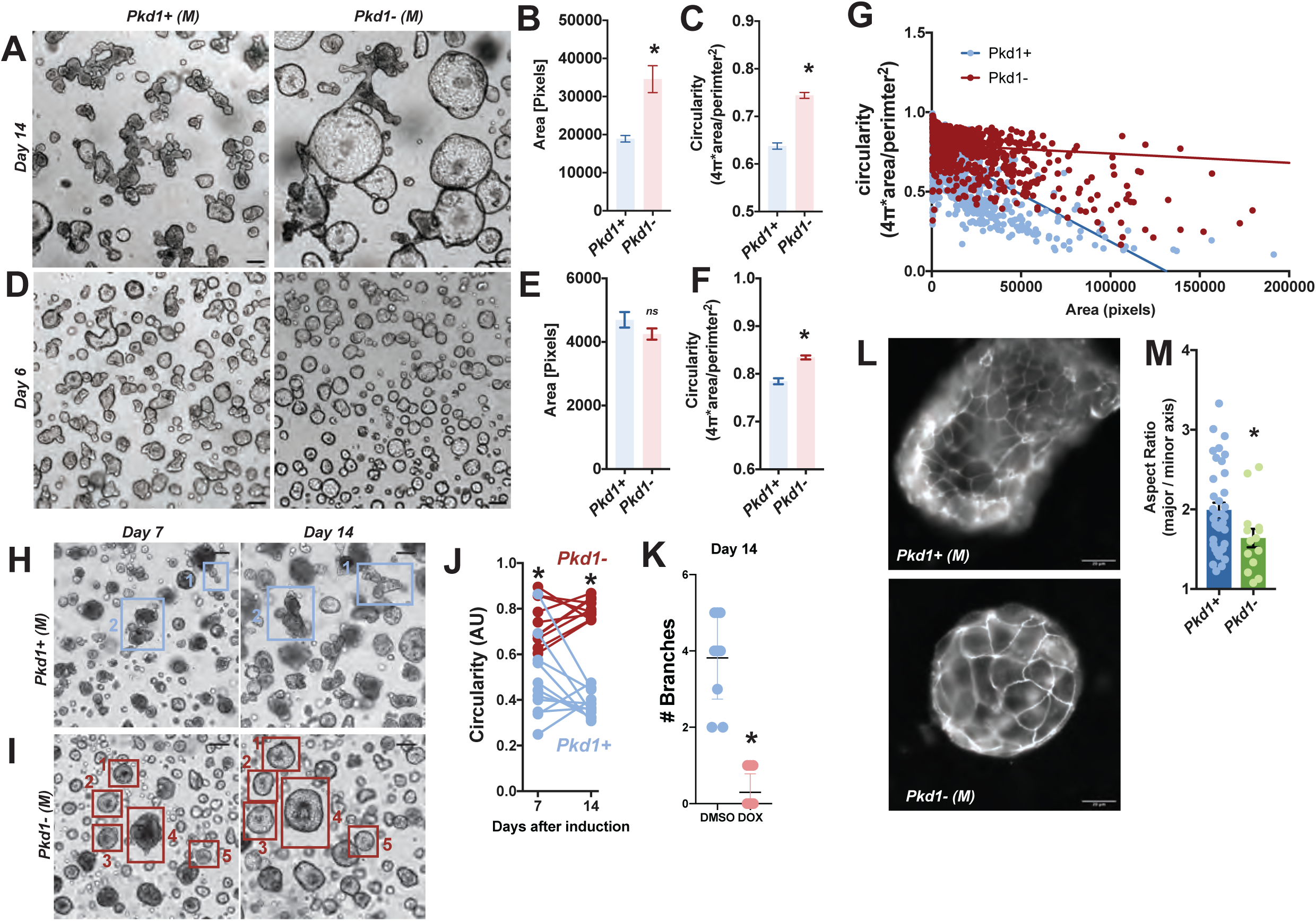
Polycystin expression regulates cell and tubular morphology in 3D cultured renal epithelial cells. (A) Immortalized mouse *Pkd1^fl/fl^ Pax8rtTA, TetO-Cre* medullary (M) renal epithelial cells, clone #313 cells treated with vehicle (DMSO = *Pkd1*+(M)) or with doxycycline (DOX = *Pkd1*- (M)) and then cultured in Matrigel for 14 days. Semi-automated quantification from three separate experiments of tubuloid size (B) and (C) circularity show significant differences in both size and circularity (*Pkd1*+(M) n= 836; *Pkd1*-(M) n=595; +/− SEM, P<0.05, two tailed Student’s T-Test). (D) The same tubuloid cultures after only 6 days in 3D culture show an early significant difference in circularity (F) after *Pkd1* knockout but not in size (E) (*Pkd1*+ (M) n=372; *Pkd1*- (M) n=452; +/− SEM, P<0.05, two tailed Student’s T-Test). (G) The relationship between size and circularity in the *Pkd1*+(M) and *Pkd1*-(M) tubuloids was analyzed with linear regression. Loss of *Pkd1* resulted in increased circularity with little correlation with size (n=2476, R2= 0.08269). In contrast, in tubuloids with normal *Pkd1* expression, there is a significant negative correlation between circularity and size: the larger the tubuloids became, the less circular they appeared (p<0.001, n= 1536, R2 = 0.4135). (H,I) In separate experiments individual tubuloid structures were tracked through time. *Pkd1* (M) (clone# 313) cells treated with (I) or without doxycycline (H) from day 7 until day 14 in 3D culture. The *Pkd1*+(M) tubuloids became less circular (J) through time with significantly more branches (K) than the *Pkd1*-(M) structures (*Pkd1*+(M) n= 11; *Pkd1*-(M) n=10; +/− SEM, P<0.05, two tailed Student’s T-Test). (L) *Pkd1* (M) (clone# 313) cells treated with or without doxycycline and cultured in Matrigel for 14 days, fixed, and labeled with junctional marker ZO1. (M) Individual cell shape (aspect ratio / AR) as visualized with ZO1 demonstrates a significant alteration in cell shape after loss of *Pkd1* expression (*Pkd1*+(M) n= 37; *Pkd1*-(M) n=14; +/− SEM, P<0.05, two tailed Student’s T-Test).

### Key cell shape protein Ezrin interacts with the polycystins

To explore how the polycystins regulate cell morphology / shape we turned to a bioinformatic approach utilizing a recent PC1 interactome data set of a proteomic study identifying proteins that interact with endogenous mouse PC1 (in the brain) created by Lin et al^22^. We compared the PC1 interactome with the GO term gene/protein list for cell shape (GO:0008360; Figure 2A) and mapped the potential functional interactions of the overlapping genes / proteins using a String analysis^24^ (Figure 2A). At the heart of the String interaction map of the genes/proteins that are both GO terms for cell shape and are part of the PC1 interactome, are the two ERM genes/ proteins *Ezr* /Ezrin and *Msn* /Moesin. Ezrin, with the highest number of demonstrated and predicted connections, binds the F-actin cytoskeleton to the apical surface via its interactions with phosphoinositides like PIP2 (PI4,5P), and is critical for apical membrane structures like the brush border and the apical compartment^25,26^.

**Figure 2:**
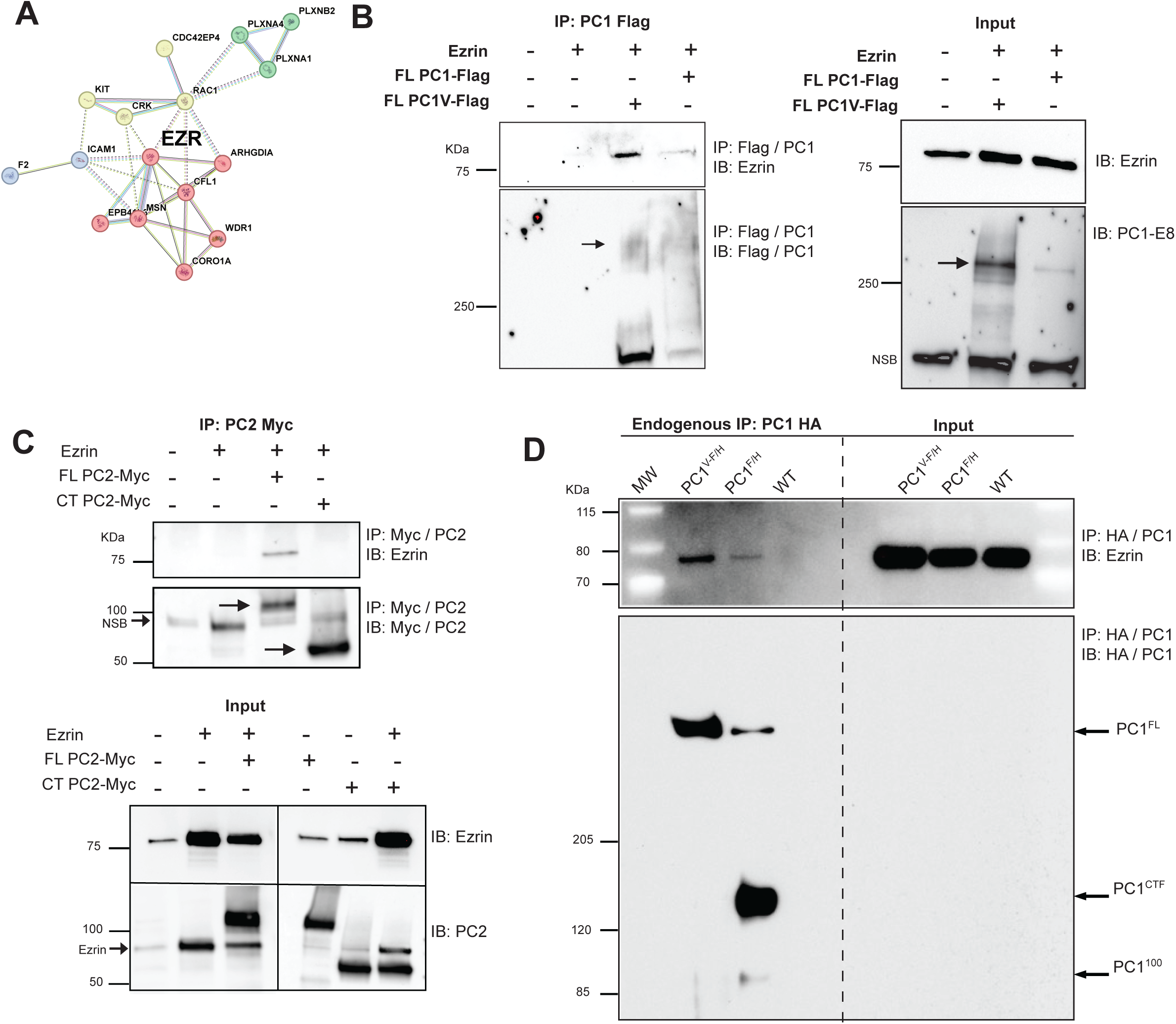
Polycystins interacts with key cell shape protein Ezrin in the kidney. (A) Bio-informatic comparison of GO terms for cell shape (GO:0008360) and the mouse PC1 brain interactome (derived from mouse brain, Lin *et al* 2023^22^). Key ERM proteins Ezrin (*EZR*) and Mosin (*MSN*) are central in the predicted interactions (String analysis). (B) Immuno-precipitation experiments using overexpressed FL-PC1 - Flag, FL-PC1V - Flag, and Ezrin constructs expressed in HEK293 cells demonstrate that the IP with Flag M2 beads can pulldown Ezrin (100μg used for IP). Flag-M2 beads pulldown product probed with the PC1-E8 antibody^15^ confirmed the presence of PC1 and PC1V. Input blots (1% for ezrin IB, 5% for PC1 IB) probed for Ezrin and PC1 confirms the presence of the Ezrin and PC1 constructs in the reaction. (C) Immuno-precipitation using over-expressed PC2-Myc and Ezrin constructs demonstrate the full length PC2 interacts with Ezrin, however the shorter C-terminal tail construct (CT PC2-Myc) does not (NSB = non-specific band). Blotting the Myc pull down with anti-Myc reveals the PC2 constructs were present in the reaction, and the inputs further confirm. (D) Endogenous PC1 (PC1^F/H^) and PC1V (PC1^V-F/H^) from mouse kidney epithelium interact with endogenous ezrin. Kidney epithelial cells were harvested at P4-P5 pups and cultured for 4 days. 2mg lysate used for IP and the input used 50μg. Immuno-precipitation of the endogenous PC1-HA was completed with HA, and the pulldown product probed for Ezrin. IB of the pulldown product demonstrates the presence of the endogenous tagged forms of PC1. Input demonstrates endogenous Ezrin is abundant in renal epithelial cells.

First we wanted to understand if Ezrin interacts with PC1 in renal epithelial cells. To test this hypothesis we began with immunoprecipitation experiments with tagged PC1 and PC2 protein and Ezrin in HEK293 cells. Flag tagged full length PC1, and PC1 with the mutation, Thr to Val, at position 3041, which prevents the autocleavage event at the GPS site necessary for trafficking of PC1 and the polycystin complex to the primary cilium (PC1V), both successfully pulled-down Ezrin (Figure 2B). The success of uncleaved PC1 in interacting with Ezrin suggest their interaction is outside the primary cilium and may be early in polycystin maturation. Next we did the same with myc-tagged PC2, again the full length PC2 successfully pulled down Ezrin, but an artificially truncated C-terminal PC2 construct does not (Figure 2C). These data support a physical interaction of the polycystin with Ezrin in a transfection system, and that cleavage of PC1 is not necessary for the interaction.

Finally, we wanted to confirm that endogenous polycystins can pull down endogenous Ezrin in renal epithelial cells. We used the *Pkd1F/H-BAC* transgenic mouse line^27^, that has three copies of a mouse *Pkd1 BAC* transgene with an NH2-terminal triple-FLAG epitope tag and a C-terminal triple-hemagglutinin (HA) epitope tag, expressed under the control of the native *Pkd1* promoter^27^ and the modified *Pkd1V-F/H-BAC*, that harbors 3-4 copies of the *Pkd1V* transgene (GPS cleavage T/V mutation at 3041 that prevents cleavage)^28^. Kidney epithelial cells were harvested from P4-P5 pups and cultured for 4 days and then used for the immunoprecipitation experiment. Endogenous PC1 and PC1V successfully pulled down endogenous Ezrin (Figure 2D). Overall, we observed relatively similar expression levels and immunoprecipitation efficiency for PC1 (full length + CTF+ P100) and PC1V (Figure 2D lower panel), but the level of Ezrin pull down was far more robust with the uncleaved PC1V, again increasing the support that Ezrin interacts with an uncleaved form of PC1, before it traffics to the primary cilium.

We next sought to better understand where Ezrin was interacting with the full length PC1 and PC2. We investigated if a common affinity for a membrane specific type of phosphoinositide may both facilitate the interaction and reveal where the polycystins and Ezrin interact in the cell. Ezrin localizes to membranes via electrostatic interaction with phosphoinositides like PI(4,5)P2^25,26^, and TRP channels (like PC2/ TRPP2) are also well known to interact with phosphoinositides in the plasma membrane^29^, could these electrostatic interactions be facilitating the Ezrin /PC1 /PC2 interactions? We used a lipid overlay assay (PIP strips) to first understand if PC2 (TRPP2) binds phosphoinositides and specifically if the species overlapped with those for Ezrin. The pattern of affinity for PC2 / Ezrin and the phospholipids were extremely similar with 7 specific common species of phosphoinositides: Ptdlns(3)P, Ptdlns(4)P, Ptdlns(5)P, Ptdlns(3,5)P_2_, Ptdlns(4,5)P_2_, Ptdlns(3,4,5)P_3_, and Phosphoatidylserine (Supplemental Figure 3). Excitingly, Xu et al^30^ previously showed that PC1, through interactions with its cytosolic PLAT domain, directly binds Ptdlns(4)P and Phosphoatidylserine, meaning all three proteins, PC1/ PC2/ Ezrin, bind these same two phosphoinositides. Ptdlns(4)P and Phosphoatidylserine are both found at the plasma membrane^31^, supporting it as a probable site of localization for the interaction between Ezrin and PC1/PC2.

### PC1 regulates expression, abundance, and localization of Ezrin in human kidneys

To explore the possible functional relationship between *EZR* and *PKD1/2* we first asked if there was evidence for a genetic interaction in human ADPKD kidneys. We compared the gene expression (mRNA levels) of *EZR* in cortical and medullary tissue samples from normal human kidneys and nephrectomized ADPKD kidneys (Supplemental Figure 2A) and found in both cortex and medulla, *EZR* gene expression is significantly increased in the ADPKD kidney samples (p<0.05). Data available from Muto et al^32^ allows finer resolution, demonstrating increased *EZR* in proximal, TAL, DCT, and CD segments (Supplemental Figure 2B) of the neprectomized end-stage ADPKD human kidneys^32^. We next measured the Ezrin protein abundance in cortical tissue samples from male normal human kidneys (NHK; n=4 unique individuals) and from cortical cysts from male ADPKD (*PKD1* variants) kidneys (Figure 3A; n=5 unique individuals) and found to our surprise the amount of Ezrin protein was significantly decreased in the ADPKD cyst samples (Figure 3B; p<0.05). To understand if localization of Ezrin was altered in addition to abundance we used immunofluorescent antibody staining of paraffin embedded sections from NHK and ADPKD kidneys and found a radically altered localization of Ezrin in the ADPKD tissue. In NHK cortical tissue sections (Figure 3 C-E, scale bar 100μm), Ezrin is prominent in proximal tubule, Na+/K+ ATPase in TAL segments, and AQP2 in collecting duct principal cells with little overlap (lack of white signal, Figure 3 C-E). Interestingly, in the collecting duct, Ezrin levels in AQP2 positive principal cells is low, but it is strong in the apical compartment of the AQP2 negative, intercalated cells of the collecting duct (White arrows D and E). The marker landscape in cystic tissue from ADPKD kidneys is significantly altered (Figure 3F-J). In the epithelial cells lining the cysts, large and small, there is prominent Na+/K+ ATPase basolateral staining and Ezrin staining, when in NHK tubules these two did not overlap. In addition, AQP2 positive cells (Figure 3G) are amongst the same cells, suggesting all three markers are present in some cyst lining cells. Ezrin and Na+/K+ ATPase significantly overlap (white signal, white arrows Figure 3G, H) along the basolateral membranes, demonstrating a totally altered Ezrin localization pattern. In tubules with a strong AQP2 signal, we confirmed Ezrin mis-localization (Figure 3I, overlap signal in white, with significant Ezrin and Na+/K+ ATPase overlap), but no alterations to AQP2 or Na+/K+ATPase (AQP2 is apical with no overlap with the Nat/K+/ ATPase, Figure 3J). These data from human kidneys demonstrate that loss of the polycystin function results in altered gene expression, protein abundance, and tubule / cellular localization of Ezrin in ADPKD kidneys.

**Figure 3:**
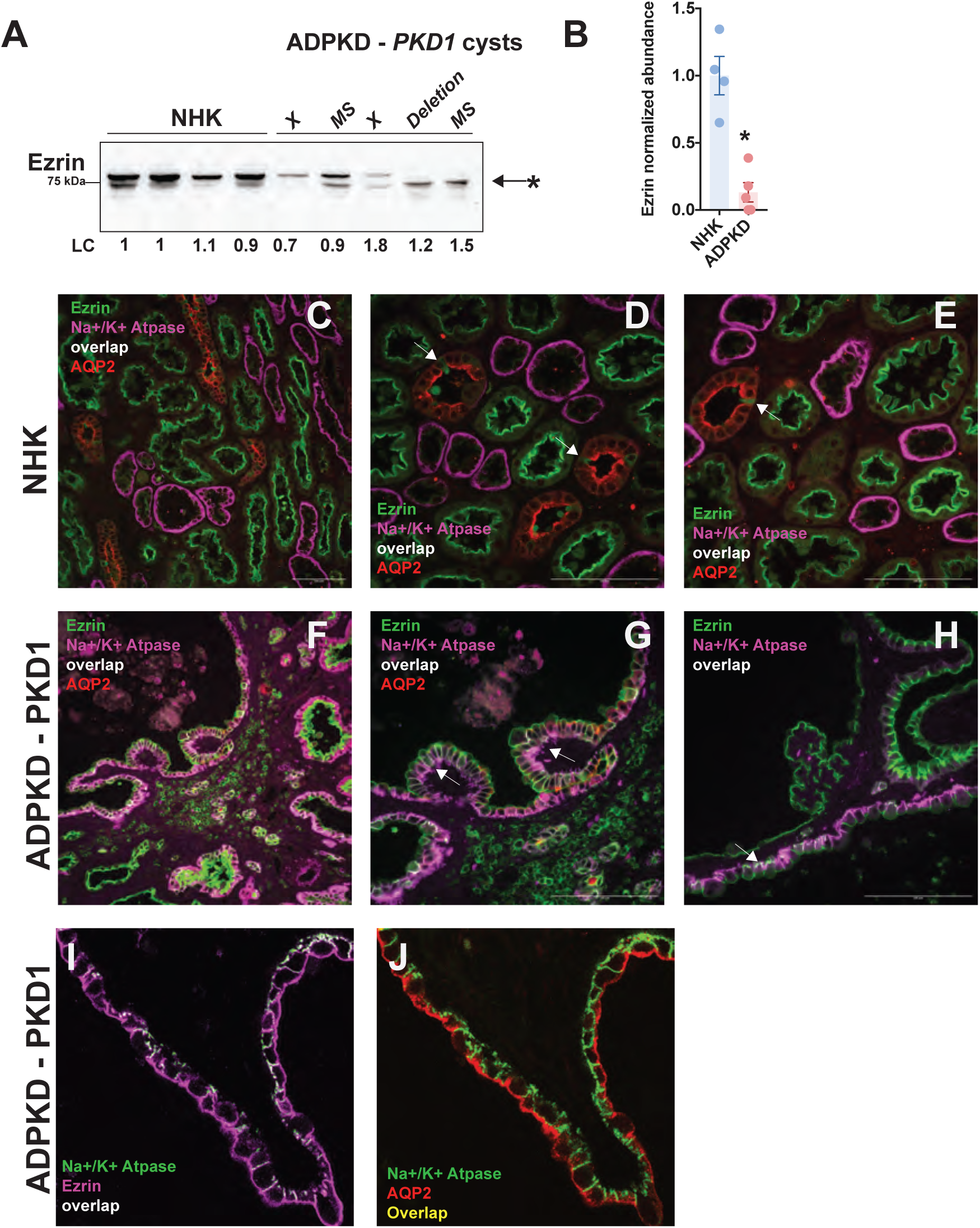
PC1 regulates abundance and localization of Ezrin in human kidneys. (A) Ezrin protein abundance in NHK kidney samples (cortex, n=4) and ADPKD samples (n=5). All samples from male donors (for ADPKD patient samples germline *PKD1* variants noted: X=truncation, MS= missense, Deletion= deletion resulting in altered reading frame). Quantified in (B) (n=4/5; +/− SE; Student T test, two tailed, p<0.05). Immunofluorescence of Ezrin (green), Na+/K+ Atpase (purple), and AQP2 (red) localization in NHK (C, 20x scale bar 100μm; D, 40x; E, 40x; overlap of ezrin and Na+/K+ Atpase is in white, there is no overlap; high Ezrin abundance in intercalated cells indicated with white arrow) and ADPKD kidney samples (F, 20x scale bar 100μm; G, 40x; H, 40x; overlap of Ezrin and Na+/K+ Atpase is in white, there is extensive overlap, sample from patient with germline *PKD1* variant c.974A>G (p.Tyr325Cys)). Ezrin localization in NHK cells is exclusive to the apical membrane (no overlap with Na+/K+ ATPase), in ADPKD tissue a significant amount of Ezrin overlaps with Na+ /K+ ATPase indicating an altered localization. (I, J) Sections of a large cyst wall (from patient with germline *PKD1* variant c.4387C>T (p.Gln1463*)) with immunofluorescence of Ezrin (purple), Na+/K+ Atpase (green), and AQP2 (red).(I) Ezrin overlaps significantly with Na+/K+ Atpase (white) demonstrating significant mis-localization of Ezrin to the basolateral membrane. (J) AQP2 is apical with no overlap with Na+/K+ Atpase in the same cyst wall. Representative images of samples imaged from n=3 NHK and n=4 ADPKD (*PKD1* variants) patients.

### Acute deletion of *Pkd1* results in significant loss of Ezrin

The epithelial cells of cysts investigated from a nephrectomized, end stage ADPKD kidney may have lost polycystin function decades earlier, and likewise are exposed to a significant kidney injury environment for a similar protracted time frame. Thus differentiating the 1^st^ order effects stemming from polycystin loss from 2^nd^ and higher order alteration from injury is difficult^23^. To help understand the relationship between Ezrin and specifically polycystin function, we next used our inducible *Pkd1* knockout cell lines. Western blot analysis of Ezrin abundance (Figure 4A) in *Pkd1+/−(C)* 2D cultured cells (cortical clone #302; Supplemental Figure 1) and *Pkd1+/−(M)*(clone #313) demonstrates that acute loss of polycystin expression after doxycycline treatment results in significant loss of Ezrin protein (Figure 4B, p<0.05, n=4 for each clone / treatment). Immunofluorescent staining with Ezrin and junctional marker ZO1 demonstrates that in the control *Pkd1+(M)* cells (Figure 3C; DMSO), Ezrin is predominantly at the apical surface, in the *Pkd1-(M)* cells (Figures 4D; DOX) the ezrin signal is significantly diminished (Figure 4E) supporting our western blot findings. When the same cell line is grown in 3D culture using our tubuloid protocol the differences in Ezrin abundance are even more stark (Figure 4F-I). The majority of cells in control tubuloids (Figure 4F,G) are positive, with a magnified view (Figure 4F white box, Figure 4G) revealing prominent Ezrin signal on the apical / luminal surface of the tubuloids. In the large spheroids that result from the loss of polycystins, a significantly reduced number of cells are positive for Ezrin (Figure 4H,I, p<0.05), again similar to what we observed in the 2D culture. The reduction in Ezrin after acute polycystin loss in 2D cultured cells or 3D tubuloids/ spheroids mirrors our observations in end stage ADPKD kidneys, suggesting polycystin control of Ezrin abundance is a proximate, or a first order interaction.

**Figure 4:**
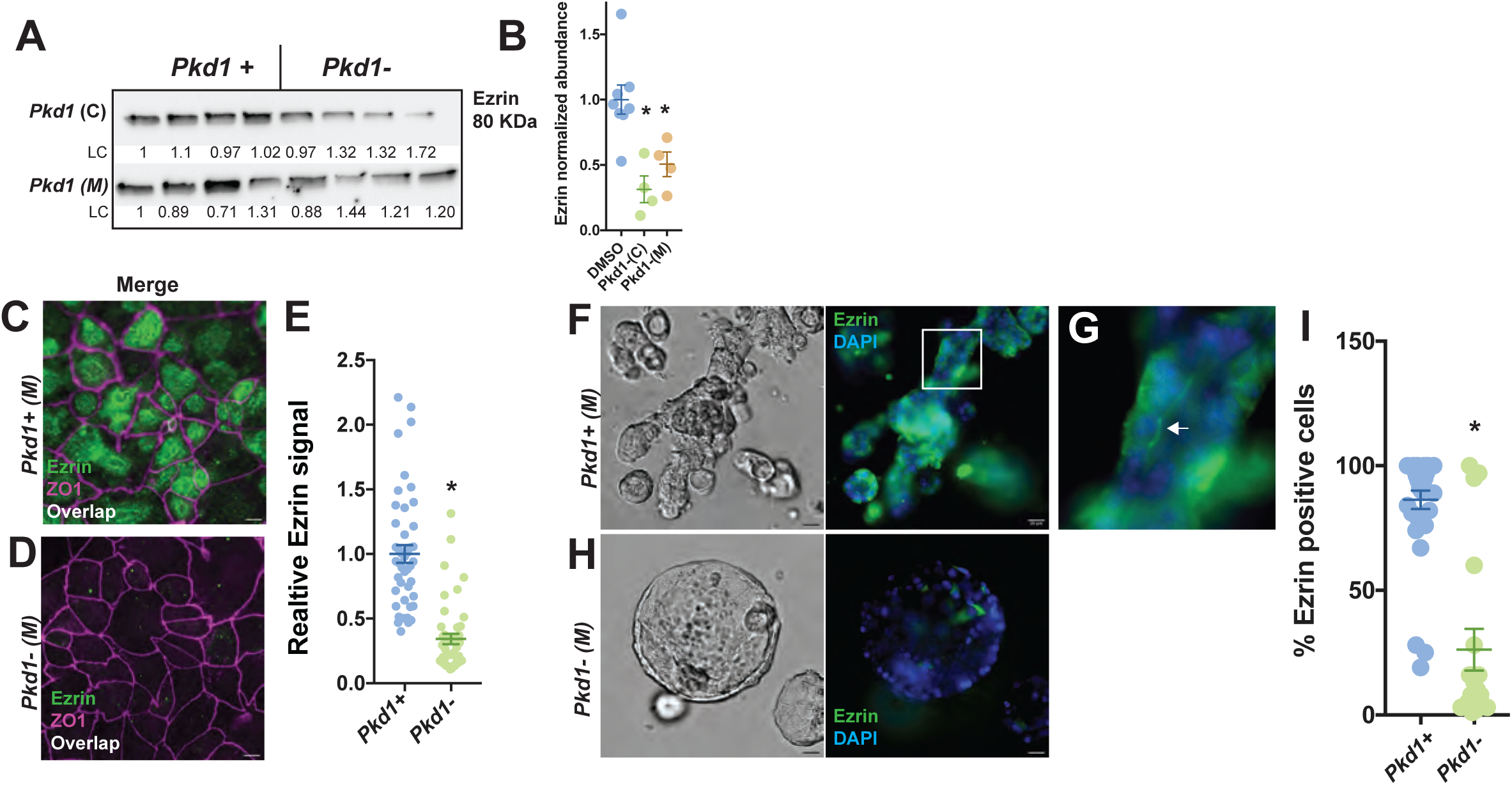
Acute deletion of *Pkd1* results in significant loss of Ezrin. (A) Western blot for Ezrin abundance in immortalized mouse *Pkd1*^fl/fl^ *Pax8rtTA, TetO-Cre* cortical (C) renal epithelial cells, clone #302 (*Pkd1*+/− (C)) and clone #313 (*Pkd1*+/− (M)), cultured with vehicle (DMSO = *Pkd1*+) or doxycycline (DOX = *Pkd1*-). LC is loading control calculated from total lane protein. (B) Quantification of Ezrin abundance in A, n=4 for each clone and each treatment group; +/− SEM, p<0.05, ANOVA with secondary Dunnett’s for multiple comparisons. (C,D) Immunofluorescent labeling of Ezrin and ZO1 in *Pkd1* (M) (clone #313) in 2D culture in control (C) or after treatment without doxycycline (D) with quantification. Ezrin localizes to apical surface and does not overlap with ZO1 in *Pkd1*+ control cells. Inducible knockout of *Pkd1* (DOX = *Pkd1*-) led to significant reduction in Ezrin signal. Exposure and Intensity matched, representative of n=4 experiments for each treatment group; n=45 cells for measured Ezrin immunofluorescence integrated intensity, +/− SEM, p<0.05, two tailed Student’s T-Test). (E-H) Tubuloids grown in 3D culture using *Pkd1* (M) cells (clone #313) treated with DMSO (E, control *Pkd1*+ (M)) or doxycycline (G, DOX, *Pkd1*- (M)) imaged in brightfield and with labeling for Ezrin (green) and DAPI (blue). (F) Close up of control (DMSO) tubuloids showing Ezrin localization on the luminal surface (white arrow). *Pkd1* knockout (*Pkd1*-) results in large hollow spheres containing few cells with Ezrin (G). (H) Quantification of 3 separate experiments (*Pkd1+(M)*(DMSO), n=34 tubuloids, *Pkd1- (m)*(DOX), n=18 tubuloids, +/− SEM, p<0.05, two tailed Student’s T-Test).

### Acute deletion of *Pkd2* results in significant loss of Ezrin *invitro* and *invivo*

Is PC1 alone or in complex with PC2 critical for Ezrin abundance? We next used the *Pkd2^fl/fl^ Pax8rtTA, TetO-Cre* (medullary clone#125; *Pkd2+/−(M)*) cell line^33^, to investigate a role for PC2 in controlling Ezrin abundance. *Pkd2+/−(M)* cells grown in 2D culture treated with or without doxycycline were lysed and Ezrin abundance evaluated using Western blot (Figure 5A). *Pkd2- (M)* cells (DOX) had significantly less Ezrin abundance (Figure 5B, n=7 for each treatment, p<0.05). Immunofluorescence using antibodies to Ezrin and ZO1 (5C,Ci, D, Di) show that in control, *Pkd2+* cells, Ezrin is apical, located in structures consistent with brush border extensions of the membrane (Figure 5Ci). *Pkd2-* cells (DOX) show altered cell morphology and reduced Ezrin, displaying a diffuse signal across the entire cytoplasm (Figure 5D,Di), similar to the reductions observed with the loss of PC1 (Figure 4). Next we used primary renal epithelial cells from *Pkd2^fl/fl^ Pax8rtTA, TetO-Cre, mTmG* mice (as described previously^14^) in 3D culture to form tubuloids. Structures from *Pkd2-* cells (treated with DOX) and control *Pkd2+* (DMSO) were retrieved from the Matrigel after 2 weeks of culture and total Ezrin and PC2 abundance was measured by Western Blot (Figure 5E). Ezrin abundance was reduced significantly and correlated with the reduction of PC2, both reduced more than 50% (Figure 5F; n=3 for each treatment, p<0.05). Taking advantage of the mTmG reporter to reveal the cell specific *Pkd2* genotype (Figure 5G, Red=*Pkd2*+; Green=*Pkd2*-; verified by cell sorting^14^) we co-stained the tubuloid / spheroid structures with Ezrin (Figure 5H, Ezrin =purple signal, Ezrin/*Pkd2*-overlap = white) to observe, in a single cell, the relationship between *Pkd2* genotype and Ezrin localization. Remarkably, Ezrin was most abundant in the *Pkd2+* cells, with very little overlap with the green, *Pkd2-* signal, as exhibited by quantified colocalization Ezrin and *Pkd2+* signal (Figure 5I, J; p<0.05). These data support a role for both polycystins in regulating Ezrin.

**Figure 5:**
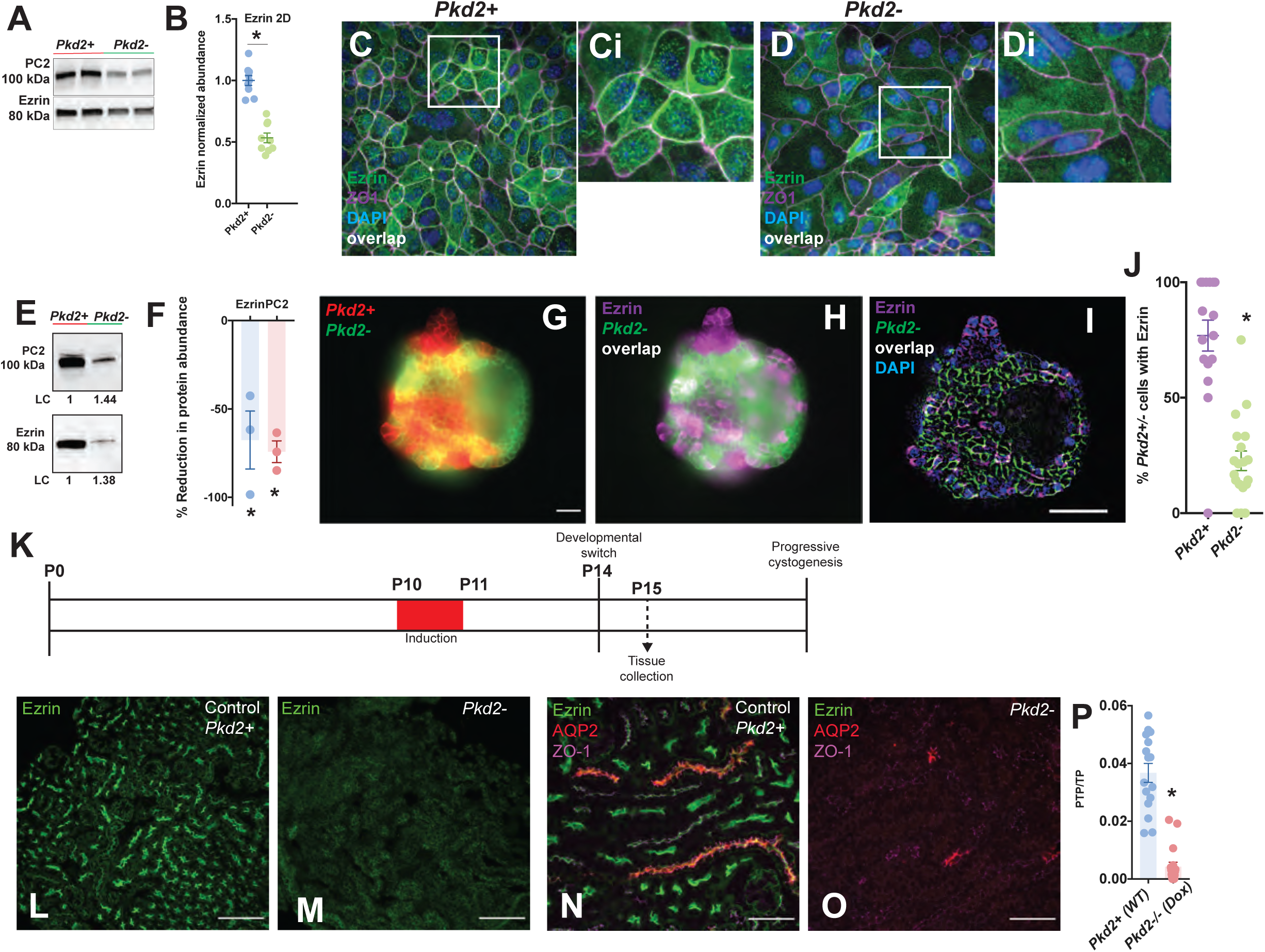
Acute deletion of *Pkd2* results in significant loss of ezrin *in vitro* and *in vivo*. (A) Mouse immortalized *Pkd2^fl/fl^ Pax8rtTA, TetO-Cre* (medullary clone#125) cells grown in 2D culture, treated with vehicle DMSO (*Pkd2+*) or doxycycline (*Pkd2-*) for 7 days and assessed for Ezrin and *Pkd2* abundance by Western blot and (B) quantified (n= 7 for each treatment, +/−SEM, P<0.05 two tailed student’s t-test). (C,D) Same *Pkd2* clone #125 cells grown in 2d culture treated with vehicle DMSO (*Pkd2+*) or doxycycline (*Pkd2-*), fixed and labeled for Ezrin (green), ZO1 (purple), and DAPI (blue), (representative of 3 unique experiments for each treatment, 40x, scale bar 20um). White box represents the enlarged area in Ci and Di. (E) Tubule fragments (from *Pkd2^fl/fl^ Pax8rtTA;TetOCre; mTmG* mice) were used to generate 3D tubuloids treated with vehicle or doxycycline, lysed, and used for western blotting to quantify PC2 and Ezrin abundance in the 3D structures. (F) Quantification showed a significant decrease in both PC2 and Ezrin abundance when compared to controls (n=3, +/ SEM, p-value <0.05, two tailed student’s t-test). (G, H) Immunofluorescent images demonstrate the mTmG (*Pkd2*+ red, *Pkd2*-green) reporter following treatment with doxycycline alongside ezrin (purple) and *Pkd2*-(green) with overlap (white). Magnification 20X, Scale bar 50μm. Representative of n=3 unique experiments for each treatment. (I,J) The colocalization quantification of ezrin and *Pkd2* signal exhibits a statistically significant decrease in *Pkd2*-cells compared to *Pkd2*+ (n=19 unique tubuloids, and 214 total cells, two-tailed p-value <0.0001). (K) *Pkd2^fl/fl^ Pax8rtTA; TetOCre* mice (P10) were treated with doxycycline for a short induction period to inactivate *Pkd2* in the kidneys^28^. Representative P15 kidneys (time point before significant cyst development) were stained from both control (L) and doxycycline (M) treated mice for ezrin (green). Magnification 20X, scale bar 100μm. (N, O) Higher magnification images of ezrin in control and doxycycline treated mice co-labeled with apical AQP2 (red) and junctional ZO1 (purple). Magnification 40X, scale bar 50μm.(C) There is a significant decrease in ezrin intensity (PTP/TP) in doxycycline treated mice (n=3 male mice, 5 field of views each) when compared to controls (n=3 male mice, 5 field of views each) (two tailed p-value<0.0001).

Finally, we investigated if the relationship between *Pkd2* expression and Ezrin abundance was also found *in vivo*. We used *Pkd2^fl/fl^ Pax8rtTA, TetO-Cre* male mice treated via the Short Induction model^34^ (Figure 5K) where doxycycline is administered on P10-P11 pups and tissue is collected at P15, before the development of large cysts^34,35^. Kidneys from animals treated with DOX or the controls were removed at P15 fixed and sectioned and stained for Ezrin (L-O). Control kidneys at P15 look normal and have high Ezrin abundance in the cortex (L, scale bar 100μm) in proximal tubule and AQP2 positive collecting ducts (Figure 4N, scale bar 50μm). In the *Pkd2-/-* animals (Figure 4M,O), tubule structure still appears normal but there is dramatic reduction of Ezrin abundance (Figure 4P, p<0.05). Interestingly, at later time points (P25) with significant cystogenesis, Ezrin abundance is reduced but not to the extent observed at P15 (Supplemental Figure 3) indicating a specific role for polycystin regulation of Ezrin early in the postnatal period.

### Polycystin loss prevents Ezrin activation and apical localization in renal epithelial cells

To further understand the PC1 /Ezrin interaction, we immunoprecipitated Ezrin in HEK293 cells using an Ezrin antibody, 3145, that specifically recognizes a conformational epitope (Figure 6A). The 3145 epitope is located in the alpha linker region of the Ezrin protein (Figure 6B) a region available only after Ezrin has gone through conformational changes facilitated by binding to phosphoinositides like PIP2 and a secondary phosphorylation event at residue T567 (Figure 6C), a sequence we here refer to as Ezrin activation. We found the antibody could only successfully bind the native Ezrin when PC1 was co-expressed (Figure 6A). The necessity of inclusion of PC1 to facilitate the binding of 3145 to Ezrin suggests that PC1 plays a role in the activation of Ezrin. We explored this possibility by combining the 3145 Ezrin antibody that preferentially binds the active Ezrin with a second Ezrin antibody 3c12^36^ with a different epitope (Figure 6B) that preferentially binds the inactive or closed confirmation of Ezrin^36^ (Figure 6C). We labeled 2D cultured *Pkd1+/−(M)*(clone #313) control cells (DMSO= *Pkd1+*) and cells treated with doxycycline (DOX = *Pkd1-*) for Ezrin with both antibodies, and ZO1 (Figure 6D-J). Control cells demonstrated that activated Ezrin localized exclusively to the apical surface, but the inactive form of Ezrin localized to the cell periphery in pattern similar to the junctional maker ZO1 (Figure 6D, E), but apical of ZO1 (Figure 6F) as demonstrated in the Z projection. A plot of relative intensities across a linear axis covering one cell (overlay in Figure 6D and Figure 6G) further illustrates the exclusive apical localization of the active Ezrin (green trace) and the similarity but not perfect overlap of the inactive Ezrin (orange arrow) and ZO1 (purple arrow). *Pkd1-* cells (DOX) show an almost complete loss of the active form of Ezrin (Figure 6H-J) with little change in the inactive form. Interestingly, the distinction between the inactive Ezrin and ZO1 localization is also lost, (Figure 6J, orange arrow) confirmed with the perfect overlap in the signal intensity profile (Figure 6K orange and purple arrows). These data strongly support a role for the polycystins in regulating the active form of Ezrin that localizes to the apical surface of renal epithelial cells.

**Figure 6:**
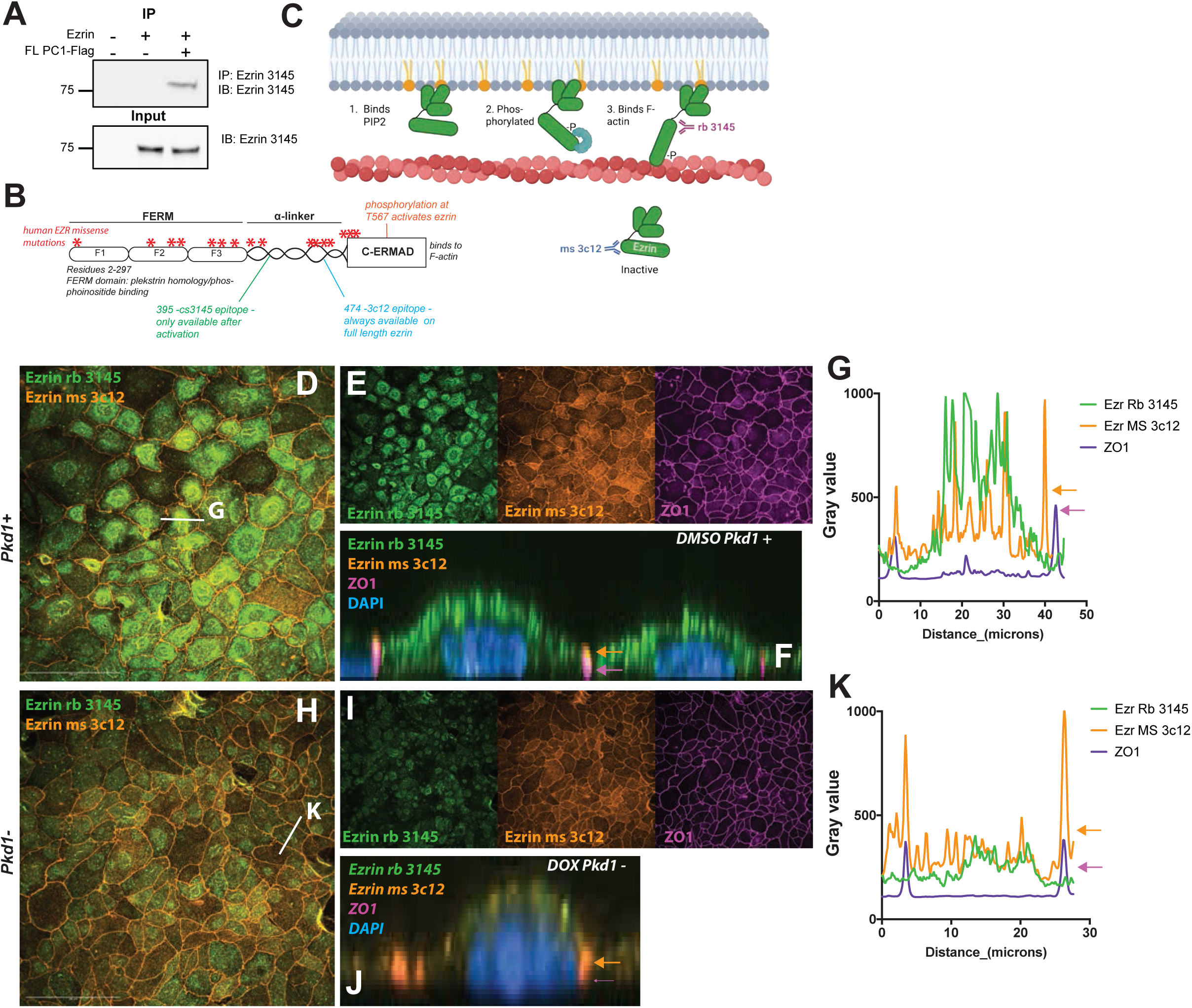
Polycystin loss prevents Ezrin activation and apical localization in renal epithelial cells. (A) Reverse IP using constructs described in Figure 5, where the Ezrin antibody 3145 is used to immunoprecipitate Ezrin unexpectedly revealed that the presence of the FL PC1-Flag construct was necessary for the Ezrin 3145 antibody to bind the over expressed Ezrin protein, suggesting PC1 may effect Ezrin folding and activation status. (B) Map of the human Ezrin protein with key domains labeled, human missense variants, activating phosphorylation site (T567), and the location of the epitopes for the Ezrin cs3145 antibody (green) and the Ezrin 3c12 antibody (blue). (C) Ezrin activation is a multistep process, beginning with Ezrin forming electrostatic interactions with Phosphoinositides like PIP2 at the apical surface, followed by phosphorylation at the T567 site, an unfolding of the protein and finally interactions with the actin cytoskeleton via the C-ERMAD domain. Previous work demonstrated that the Ezrin 3145 protein can only bind a natively folded ezrin protein that has been “opened” via phosphorylation^36^. Here we use the Ezrin 3145 antibody to label activated ezrin and the Ezrin 3c12 antibody to label the inactive form. (D-K) Immortalized mouse *Pkd1*^fl/fl^ *Pax8rtTA, TetO-Cre* renal epithelial cells, clone #313 (*Pkd1*(M)), cultured with vehicle (D,E,F; DMSO = *Pkd1*+) or doxycycline (H, I,J; DOX = *Pkd1*-) labeled with Ezrin 3145 (active form) and Ezrin 3c12 (inactive). D, H merged image of the two Ezrin signals demonstrate that in control *Pkd1*+ cells (DMSO) the two Ezrin forms do not overlap, Ezrin 3145 signal is apical (green) and the Ezrin 3c12 localizes to the cell edges (orange), in the *Pkd1*-cells (DOX) there is loss of the Ezrin 3145 signal, but not the Ezrin 3c12. Scale bar 100μm. E, I images of individual wavelengths, Ezrin 3145 (green), Ezrin 3c12 (orange), and ZO1 (purple) illustrating the differences in localization, and the overlap of Ezrin 3c12 with ZO1 at the cell junctions, Representative of n=3 unique experiments for each treatment. (F, J) Z projection of the cells in D and H respectively. In the *Pkd1*+ (DMSO) cells (F), Ezrin 3145 is exclusively at the apical compartment, with the Ezrin 3c12 localized to the junctions, but distinct and above the ZO1 signals (orange and purple arrows). In (J) the *Pkd1*- (DOX) cells have lost most of the Ezrin 3145 signal and the Ezrin 3c12 overlaps the ZO1 signal. (G, K) Signal intensity (gray value) along the projected white line across one cell in D and H respectively. The traces illustrate the localization of Ezrin 3145 to the apical compartment and Ezrin 3c12 to the junctions. (G) The peak values for the Ezrin 3c12 and ZO1 are not overlapping, though close (orange and purple arrows) in the *Pkd1*+ (DMSO) cells. (K) In the *Pkd1*- (DOX) cells the green signal is greatly diminished and the Ezrin 3c12 and ZO1 peak signals completely overlap.

### Ezrin phosphorylation inhibitor phenocopies altered tubular morphology observed with polycystin knockout

The loss of polycystin expression and function significantly reduces the active form of Ezrin during cystogenesis, but is the loss of Ezrin alone also cystogenic? The three human ERM proteins, Ezrin, Radixin, and Moesin are a product of a gene duplication event unique to mammals^37^. Evidence suggests significant overlap in expression and function, with each able to compensate the loss of each other^38^. All three are expressed in the human kidney (Supplemental Figure 2) so instead of using an Ezr/Rdx/Msn triple knockout to confirm a role of Ezrin/ERMs in cystogenesis we instead used a small molecule inhibitor (NSC668394) that binds all three ERM proteins and prevents their phosphorylation at the conserved T567 residue of the C-ERMAD domain^39,40^. Because our data suggested PC1 plays a role in the activation of Ezrin we hypothesized that a small molecule that binds the phosphorylation site may interfere with the physical interaction between Ezrin and PC1. We performed further immunoprecipitation experiments, pulling down PC1 with a flag antibody and probing for Ezrin with or without the Ezrin inhibitor and found the treatment of cells with the NSC668394 inhibitor prevented the pulldown of Ezrin and the interaction (Figure 7A). Next, we turned to our 3D culture to next ask if inhibition of Ezrin activation would lead to cystogenesis. *Pkd1+(M)* (clone #313) cells treated with two different concentrations (10μM, 20μM) of NSC668394 produced distinctly spherical tubuloids similar to those observed to tubuloids generated from DOX treatment (Figure 7B). The circularity of the tubuloid/spheroids produced from the NSC668394 treatment were significantly greater than the control (DMSO) structures, and even more circular than DOX tubuloids /spheroids (Figure 7C, p<0.05). Interestingly, the size of the spheroids resulting from the NSC668394 treatment was significantly smaller than the control (DMSO) tubuloids (Figure 7D) suggesting the inhibition of Ezrin may uncouple the morphology (*circularity*) from cell proliferation and tubuloid growth (*area*). The relationship between circularity and area for the tubuloids treated with NSC668394 was distinct from either the DMSO tubuloids or the DOX (Figure 7E) again suggesting an alteration in one factor (circularity) without the other (area). Inhibition of Ezrin (and potentially the other ERM proteins) disrupts the PC1 / Ezrin interaction and partially phenocopies the cystogenesis observed after acute *Pkd1* knockout.

**Figure 7:**
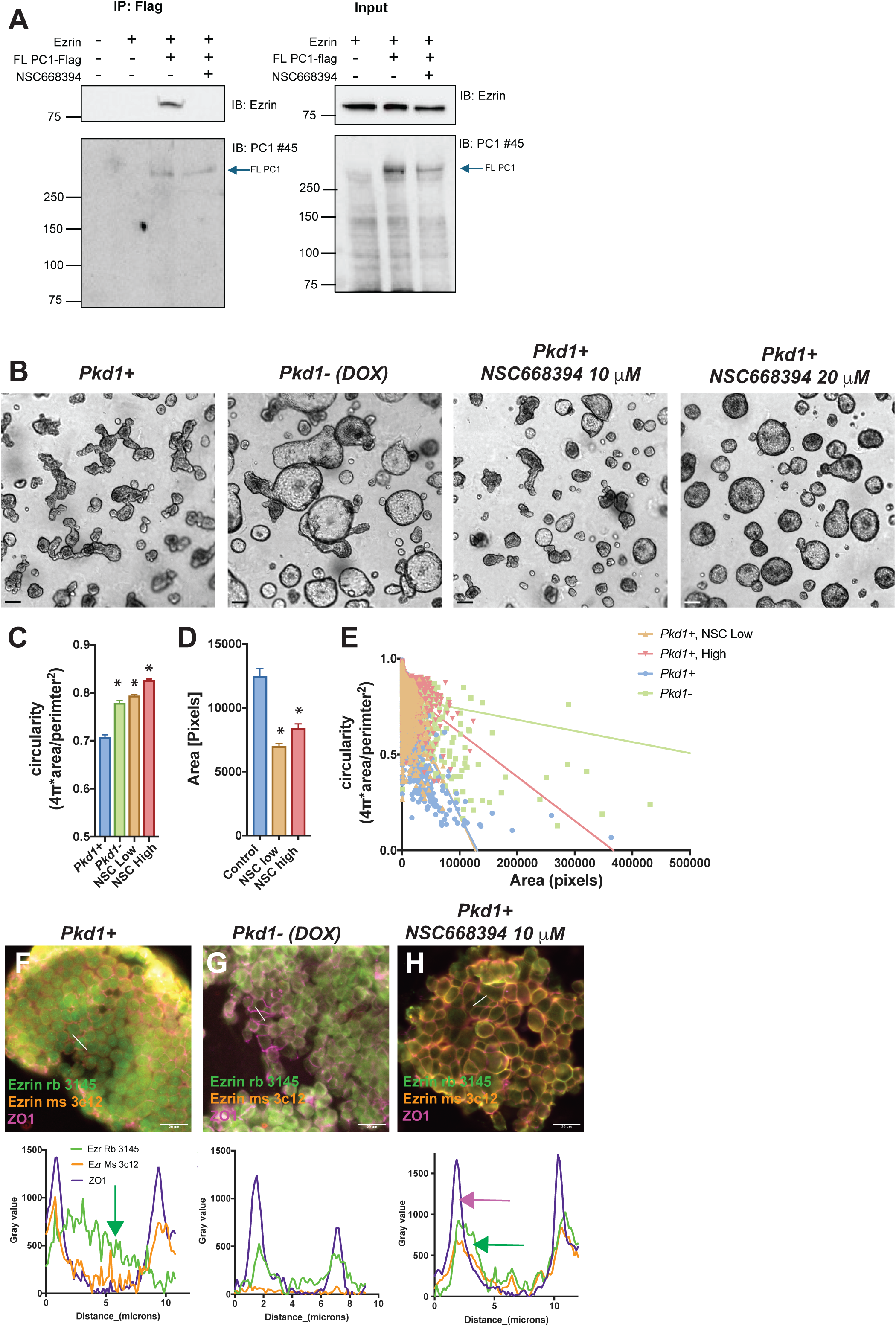
Ezrin phosphorylation inhibitor phenocopies altered cell and tubular morphology observed with polycystin knockout. (A) Immuno-precipitation experiments using overexpressed PC1 – Flag and Ezrin constructs expressed in HEK293 cells treated with or without the specific ERM phosphorylation inhibitor NSC668394 (NSC), demonstrate that the IP with Flag M2 beads can pulldown Ezrin (100μg used for IP) but not if the cells are treated for 6 hours with NSC668394 (10μM). Flag-M2 beads pulldown product probed with the PC1#45 antibody (gift from F. Qian) confirmed the presence of PC1 in the immunoprecipitation. Input blots (1% for Ezrin IB, 5% for PC1 IB) probed for ezrin and PC1 confirms the presence of the Ezrin and PC1 proteins in the lysates. (B) Immortalized mouse *Pkd1*^fl/fl^ Pax8rtTA, TetO-Cre medullary (M) renal epithelial cells = *Pkd1*(M) (clone #313) treated with or without doxycycline and then cultured in Matrigel for 14 days, with additional cultures of wildtype *Pkd1*+ (M) cells treated with two different concentrations (10μM, 20μM) of NSC668394. Semi-automated quantification from three separate experiments of tubuloid circularity (C) and (D) size (DMSO=*Pkd1*+(M) n=1391; DOX=*Pkd1*-(M) n=868; NSC Low = *Pkd1*+(M)+NSC 10μM n=1997; NSC High= *Pkd1*+(M)+NSC 20μM n=1581) found significant differences in circularity in all three treatment groups, however NSC treatment did not increase tubuloid size as observed with DOX/ *Pkd1* knockout; +/− SEM, P<0.05, ANOVA with secondary Tukey’s multiple comparison test. (E) The relationship between size and circularity in the *Pkd1*+(M), *Pkd1*-(M), and *Pkd1*+(M) treated with low and high NSC tubuloids was analyzed with linear regression. Higher doses of NSC shifted the relationship between circularity and size from tubuloids similar to control *Pkd1*+(M) tubuloids to the DOX *Pkd1*-(M) spheroids, but with a significantly different slope (p<0.05) reflective in the large difference in tubuloid size (DMSO=*Pkd1*+(M) n=1536, R2=0.4135; DOX=Pkd1-(M) n=2467, R2=0.08269; NSC Low = *Pkd1*+(M)+NSC 10μM n=4804, R2=0.1919; NSC High= *Pkd1*+(M)+NSC 20μM n=6769, R2=0.08853). (F-H) 14 day tubuloids as described above labeled with the two Ezrin antibodies 3145 and 3c12, and ZO1. (H) Like 2D cells, in the *Pkd1*+ (M) cells (DMSO) activated ezrin (green 3145 signal) is found in the cell center on the apical surface, the inactive form (orange 3c12) is found near the junction as marked by ZO1 (purple). *Pkd1*-(M) (DOX) and *PKD1*+(M) +NSC668394 (10μM) treated tubuloids both show the same loss of apical activated Ezrin (3145) and the increase of active Ezrin signal and the junctions, overlapping with ZO1. Representative images of three 3 separate experiments for each treatment (Scale bar 20μm). Signal intensity (gray value) along the projected white line across one cell in F-H respectively. The traces illustrate the relative localization of activated Ezrin (green 3145) as compared inactive Ezrin (orange, 3c12) and junctional marker ZO1 (purple).

Next we wanted to compare the effects of NSC668394 to PC1 knockout on the 3D localization of active and inactive Ezrin. We again cultured *Pkd1+(M)* (clone #313) cells, *Pkd1- (M)* (DOX), and *Pkd1+(M)* cells treated with 10μM NSC668394 in our 14 day tubuloid model, then fixed and labeled the resulting structures with Ezrin 3145, Ezrin 3c12, and ZO-1 (Figure 7F-H). Much like we observed in 2D culture, active Ezrin is found in the middle of the cell (consistent with apical localization) and away from the cell borders (Figure 7F), with the inactive Ezrin found predominantly at the cell edges, again overlapping with ZO-1. In the *Pkd1-(M)*, doxycycline treated structures, there is a profound reduction of active Ezrin in the cell center (Figure 7G). The NSC668394 treated structures also presented with a complete loss of active Ezrin in the cell center (Figure 7H), phenocopying the alterations observed in the *Pkd1-(M)* structures. The uncoupling of circularity and size observed with the Ezrin inhibitor suggests cystogenesis in our 3D model resulting from PC1 loss involves altered morphology and accelerated growth, and only morphology is related to Ezrin function.

### Inhibition of PKC activity partially phenocopies polycystin knockout and Ezrin inhibition

Inhibition of Ezrin phosphorylation recapitulates portions of the cystogenic pathway and prevents the physical interaction with PC1. Given the low likelihood of PC1 acting as a kinase, we hypothesized PC1 and or the polycystin complex may facilitate Ezrin phosphorylation by anchoring a complex containing both Ezrin and its activating kinase. The T567 phosphorylation site is the target of multiple kinases including isoforms of PKC^41^. Therefore we used a pan-PKC inhibitor compound, GO6983, to test if inhibiting PKC would recreate portions of the cystogenic phenotype. 3D culture of *Pkd1+(M)* cells (clone #313) treated with pan PKC inhibitor (PKC-inhib, G06983) (Figure 8A) showed significantly increased circularity as compared to the controls after 14 days (Figure 8B, p<0.05). In a subset of the largest tubuloids, we found a significant difference in the circularity (Figure 8D, p<0.05) of the PKC inhibitor treated tubuloids/spheroids with little change in the differences in tubuloid area (Figure 8E). We used a linear regression analysis to again quantify the relationship between circularity and area and found that the PKC inhibitor abolished the relationship between circularity and area (Figure 8F, p<0.1517). These findings are similar to what we observe in the *Pkd1-* spheroids /cysts.

**Figure 8:**
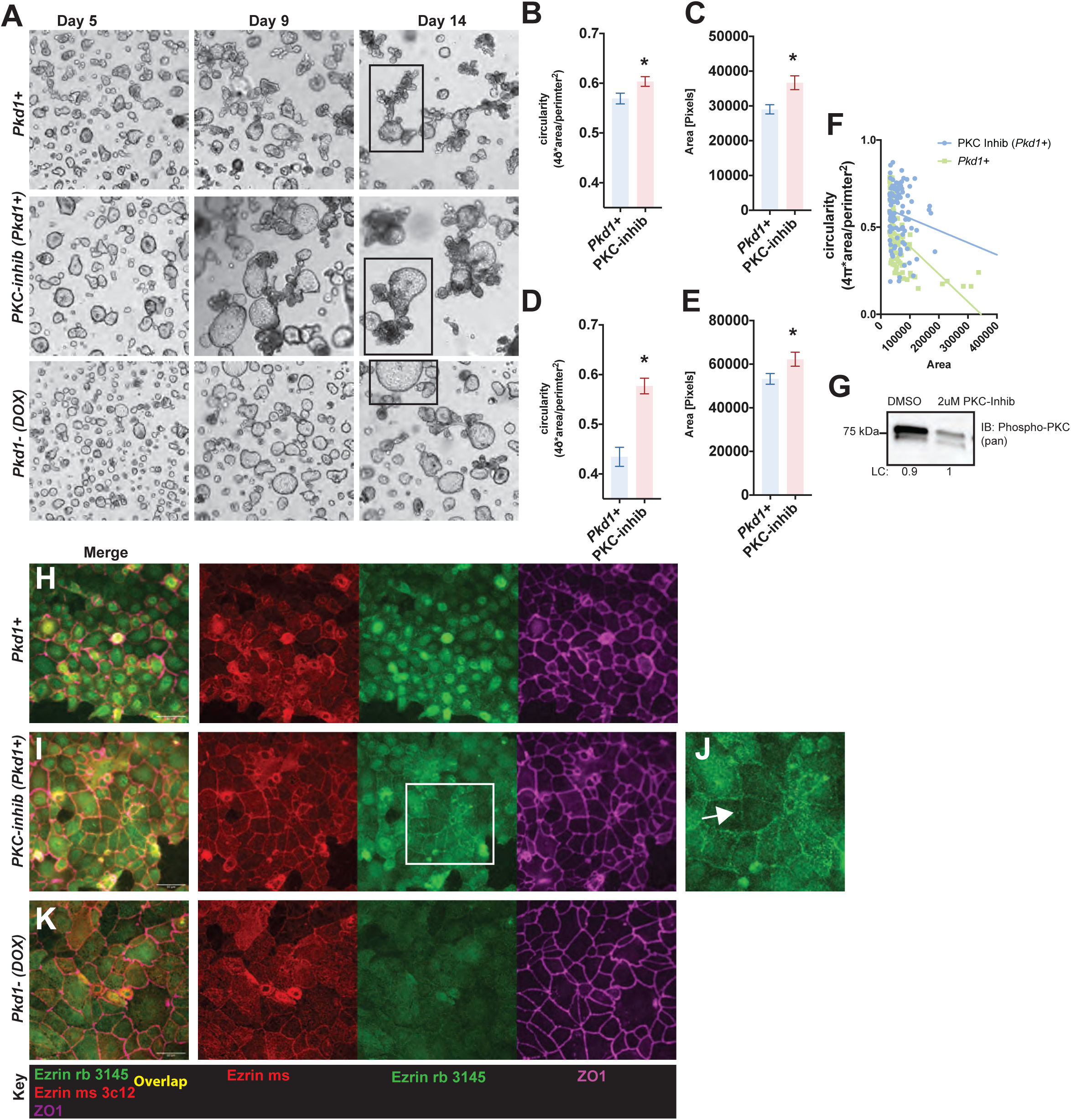
Inhibition of PKC activity partially phenocopies polycystin knockout and Ezrin inhibition. (A) Immortalized mouse *Pkd1*^fl/fl^ *Pax8rtTA, TetO-Cre* medullary renal epithelial cells = *Pkd1*(M) (clone #313) treated with or without doxycycline then cultured in Matrigel for 14 days, with additional cultures of wildtype *Pkd1*+ (M) cells treated with 2μM of pan PKC inhibitor GO6983. (B-E) Semi-automated quantification from three separate experiments of tubuloid circularity (B) and (C) size (DMSO=*Pkd1*+(M) n=273; PKC-inhib = *Pkd1*+(M)+ 2μM GO6983 n=242) found significant differences in circularity and area in the structures treated with GO6983; +/− SEM, P<0.05, two tailed Student’s T-Test. (D,E) Comparison of the largest structures (e.g. structures indicated with black box) reveals a further enhanced difference in circularity (D) for the structures treated with GO6983; DMSO=*Pkd1*+(M) n=88; PKC-inhib = *Pkd1*+(M)+ 2μM GO6983 n=103; +/− SEM, P<0.05, two tailed Student’s T-Test. (F) The relationship between size and circularity in the largest structures (*Pkd1*+(M) and *Pkd1*+(M) treated with 2μM GO6983), were analyzed with linear regression. Treatment with PKC-inhibitor abolished the relationship between circularity and size (PKC-inhib = *Pkd1*+(M)+ 2μM GO6983 n=103; p<0.1517, R2=0.02024) from tubuloids similar to what was previously observed after Doxycycline treatment (Figure 1) and treatment with Ezrin inhibitor NSC (Figure 7) supporting a common targeted pathway. (G) At the completion of the 14 day experiment structures were removed from the Matrigel and Western blot was performed to confirm the effectiveness of the GO6983 to reduce the amount of phosphorylated PKC. Blot representative of three separate experiments confirmed a significant decrease in the levels of phospho-PKC in the 3D structures after treatment with GO6983. L.C. = loading control. (H-K) Representative merged images or separated montage of Immortalized mouse *Pkd1*^fl/fl^ *Pax8rtTA, TetO-Cre* renal epithelial cells, clone #313 (*Pkd1*(M)), cultured with vehicle (H, DMSO = *Pkd1*+), doxycycline (K; DOX = *Pkd1*-), or 2μM GO6983 (I,J; PKC-inhib = *Pkd1*+ 2μM GO6983) labeled with Ezrin 3145 (active form, green), Ezrin 3c12 (inactive, red), and ZO1 (purple). Repeated in three separate experiments, scale bar 50μm. Treatment with the PKC-inhibitor reduces apical active ezrin like treatment with DOX, but unlike DOX, the PKC-inhibitor causes accumulation of the active Ezrin near the junctions (K, inset, enlarged in J, white arrow), mirroring the inactive Ezrin and ZO1 localizations.

If the polycystins facilitate the interaction between a kinase and Ezrin to regulate Ezrin activation, inhibition of the kinase should cause similar alterations in the abundance and localization of the active form of Ezrin. To test this hypothesis, we cultured *Pkd1+(M)* cells (clone #313), treated with or without the PKC inhibitor compound GO6983 (*PKC-inhib Pkd1+),* then fixed and labeled the active and inactive forms of Ezrin along with ZO1 (Figure 8H-K). Inhibition of PKC lead to a stark loss of apically localized active Ezrin (Figure 8I) with little effect on the localization of the inactive Ezrin localized to the cell periphery. Interestingly, the PKC inhibitor also appears to localize a portion of the remaining active Ezrin signal to the same cell periphery (inset Figure 8I and J, white arrow). Inhibition of PKC isoforms partially phenocopies both the inhibition of Ezrin directly, as well as the loss of *Pkd1* expression / PC1 function, supporting the hypothesis that the polycystins, outside of the cilium, facilitate a complex that includes Ezrin and its kinase, regulating Ezrin function and apical cellular morphology.

## Discussion

Here we present evidence that PC1/PC2 functions outside the primary cilium, to regulate, through molecular interactions, the important actin binding membrane tether protein Ezrin. These novel data further the hypothesis that full length PC1 functions during tubulogenesis to regulate cell and tubule shape, independent of a role in gene transcription and cell proliferation. Here we provide evidence that the 3 primary changes underlying cystogenesis: changes in cell / tubule shape, changes in cell proliferation, and changes in vectoral transport can be temporally separated. Specifically, through manipulation of Ezrin or PKC we can alter cell and tubular shape but not tubuloid size, a phenotype that in our *in vitro* system involves both cell proliferation and fluid secretion into the tubuloid /spheroid lumen. Disruption of one cystogenesis pathway, cell shape, does not necessarily proceed to the others, suggesting there is at least two separate aspects of the cystogenic pathways being regulated by the PC1/PC2 complex in our 3D invitro model. A strength of our 3D culture model choice is that it represents a reductionist approach, it contains only clonal renal epithelial cells in a static matrix environment, specifically to study cell autonomous mechanisms of cystogenesis. These characteristics, by design, reduce variability, contributing to the easily defined morphological phenotypes, but they also limit the reach of our conclusions. Our system does not model an adult functioning kidney with flow, instead it provides further insights into the role of the PC1/PC2 complex in tubule development. In this context, we conclude that the extra-ciliary, uncleaved PC1, is the likely regulator of Ezrin and cell shape primarily during tubule development, but potentially also after renal injury when developmental pathways are re-activated.

Previous work has demonstrated that uncleavable PC1 (*Pkd1*V/V) is sufficient to rescue normal kidney development, with a significant delay in the cyst development as compared to *Pkd1* knockout models^16,18^. Critically, the uncleavable PC1 is also sufficient to fully rescue the vascular and lymphatic phenotypes^21^, suggesting a conserved function of the uncleaved PC1 in tubule development across tissue types. Here we demonstrate that Ezrin is critical in PC1/PC2 regulation in kidney tubuloid shape, but the regulation of other ERM (Ezrin (*EZR*), Radixin (*RDX*), and Moesin (*MSN*)) proteins may be more important in other tissue types. Phosphorylation of the ERM protein Moesin, is critical for angiogenesis^42^ potentially via its regulation of VE-cadherin internalization^43^. Though possibly less important than Moesin, previous studies showed that knockdown of Ezrin expression in endothelial cells resulted in morphological changes, cytoskeletal reorganization^44^, and altered migration^45^ as well. These studies in endothelial cells present a similar picture to what we propose in the kidney; that uncleaved PC1/ PC2 regulate the activation / phosphorylation and localization of ERM proteins preventing disruptions in cell and tubule morphology observed when ERM protein function is lost. We predict these functions are independent of PC1/PC2 complex function in primary cilia.

We found that loss of PC1 or PC2 alters the activation and localization of Ezrin, with a significant loss at the apical surface and shift toward the cell perimeter, shifts commensurate with the observed change in cell shape. We observe similar alterations in cell morphology with the application of NSC668394, an inhibitor of ERM phosphorylation at the conserved T567, which prevents the interaction of Ezrin and actin^39^. Interestingly, Ezrin does not directly regulate the actomyosin contraction machinery, so how does it have such a profound effect on cell shape in renal epithelial cells? Previous work showed blockade of Ezrin phosphorylation at T567 (via NSC668394), depletion of Ezrin via siRNA, and masking PIP2 availability on the enter leaflet with neomycin, all sever the actin cytoskeleton from the apical surface resulting in reduced apical membrane tension, shorter and wider cells, and increased cortical tension in MDCK cells^26^. Bruckner *et al* observed after each of these Ezrin disruptions the same alteration in Ezrin location, from apical surface to the cell borders, that we observe with the loss of PC1^26^. In short, loss of Ezrin /ERMs at the apical surface shortens and broadens renal epithelial cells, the same phenomenology observed during tubule dilation and cystogenesis in ADPKD.

A role for Ezrin in cystogenesis has been considered previously. ERM proteins, Ezrin, Radixin, and Moesin, are products of paralogous genes with high amino acid and functional conservation^41^. The role of ERM proteins in regulating the apical cytoskeleton and epithelial tubular morphology was first demonstrated in model organisms that have only one ERM ortholog, for example *C. elegans* where *ERM-1* knockout results in severe morphological phenotypes including cyst formation in their nephridia epithelium^38^. In Zebrafish, the sole ERM is Ezrin which is highly expressed in the larval pronephros, and Ezrin knockout causes renal cysts^46^. Over-expression of Ezrin in Zebrafish can rescue a Joubert’s Syndrome renal cystic phenotype caused by knockout of *INPP5E*, a PIP_2_ regulator^47,48^. A specific role for ERMs in the mammalian kidney has been more complicated to dissect because all three ERM proteins are expressed in the mouse and human nephron. Ezrin (*EZR*) null mice do not survive to weaning making a renal phenotype challenging to study^49^. Segment specific mapping of ERM expression in the human kidney (based on single nuclei RNAseq experiments) shows that the greatest redundancy among the three ERMs appears to occur in the Loop of Henle while the lowest appears to be in the collecting duct where *EZR* expression is significantly higher than *RDX* or *MSN*^32^. These differential patterns of expression suggest that the collecting duct may be more susceptible to altered morphology in situations of Ezrin dysregulation compared with other segments, which is relevant for ADPKD since the collecting duct is an important site for cyst formation.

We found that in ADPKD kidneys, loss of the polycystins leads to a significant increase in EZR gene expression, but the protein levels are significantly decreased. These observations are consistent with a role for PC1/PC2 directly regulating Ezrin protein, with a loss of the positive PC1/PC2 signal leading to reduced activation and degradation of the Ezrin protein, an event that may upregulate a transcriptional response bent at correcting the loss. Understanding which transcription factors are involved in this compensatory response and whether the primary cilium is playing a significant signaling role may provide a greater understanding in how renal epithelial cells monitor their morphology in response to external pressures and signals.

Is Ezrin a cystic gene? Alterations in cell shape and morphology underlie cystogenesis but also occur in other renal injuries. Ischemia-reperfusion damages the proximal tubule and is characterized by changes in cell shape and tubule dilation. Recent work has highlighted the association of these injury phenotypes with altered cytoskeletal proteins including the loss of Ezrin^50^. Therefore loss of cytoskeletal proteins and ERMs may alter tubule shape/size but not necessarily result in cysts, which may require disruption of other signaling pathways. Here we hypothesize that after functional polycystin loss, dysregulation of Ezrin plays a critical role in the morphological changes that occur in the early stages of cystogenesis. Although *EZR* may not be a “cystic gene” further understanding of how it is regulated by the polycystins will provide new and important information about the physiological role of the polycystins and ultimately about the early pathology of ADPKD.

## Methods

### Mice

All animal experimental procedures were conducted in accordance with the University of Maryland Animal Care and Use Committee (IACUC) guidelines and procedures. The *Pkd2^fl/fl^ Pax8rtTA, TetOCre, +mTmG*^14^ *and Pkd1^fl/fl^ Pax8 rtTA TetOCre*^8^ mice were provided by the Maryland PKD center as part of the NIDDK’s Polycystic Kidney Disease Research Resource Consortium (PKD RRC). The animals used for the exvivo immunoprecipitation experiments were described previously: The *Pkd1F/H-BAC* transgenic line, which fully rescues *Pkd1* phenotypes, was a gift of Stefan Somlo^27^ (to T. Watnick), and has an NH2-terminal triple-FLAG epitope tag and a C-terminal triple-hemagglutinin (HA) epitope tag, and expressed under the control of the native *Pkd1* promoter. The *Pkd1V-F/H-BAC* transgenic mice is the same but with mutated ACT (Thr) to GTT (Val) at amino acid 3041, which abolishes *Pkd1* GPS cleavage^21^.

### Mouse Cell lines and 2D culture

Immortalized mouse *Pkd1^fl/fl^ Pax8rtTA, TetO-Cre* renal epithelial cells, clones #313 and #302 were developed by the Woodward lab and are available from the PKD-RRC. We also obtained the previously developed Immortalized mouse *Pkd2^fl/fl^ Pax8rtTA, TetO-Cre* medullary renal epithelial cells, clone #125^33^ from the PKD RRC. Briefly, these clonal cell lines were developed from cells derived from cortical or medullary regions of post-natal kidneys (P7-P10) from offspring of *Pkd1*^fl/fl^ *or Pkd2*^fl/fl^ *Pax8*-*rtTA TetO-Cre* mice^51,52^ crossed to “immorto mice” harboring a temperature-sensitive SV-40 large T antigen transgene^53^. Individual clones were screened specifically for their propensity for growth in 3D culture. Efficacy of gene deletion was determined by western blot (Supplemental figure 1). For 2D culture clonal cell lines were cultured in T75 flasks at 33°C. until confluency and fed with the Maryland PKD renal epithelia cell media (REC): 1:1 mixture of RenaLife Complete Medium (Lifeline Cell Technology LL-0025) [or Lonza REBM (CC-3191) with the REGM Single Quots supplement kit (CC-4127)] and Advanced MEM medium (Fisher Scientific #12492) with 5% FBS (ThermoFisher #26140-079), 2.2% Pen/Strep (Fisher Scientific #30-002-Cl), 0.6% L-alanyl-Glutamine (Gemini Bio-products #400-106), and 0.03% Gentamicin (Quality Biological #120-098-661). For propagation at 33°C, 10 ng/mL interferon-gamma (CST 39127) is also added to the media. Confluent flasks were then split into two, and moved to 37°C and the propagation media replaced with REC media without interferon-gamma. When cells reached 50% confluence, the cultures were treated with either 10 μg/mL of doxycycline (Sigma D3072) to inactivate *Pkd1* / *Pkd2* or DMSO for controls for 6 days. The cells were then plated on glass, grown to confluence and imaged for 2D experiments or used of the 3D experiments described below.

### Human and mouse primary cell culture and samples

Normal human kidney renal epithelial cells and tissue samples and ADPKD cyst epithelial cells and tissue samples were received from the Maryland PKD Research and Clinical Core Center (reviewed by the UMB IRB and determined to not be human research, requiring no further IRB review), as part of the PKD-RRC. Primary cells were cultured on plastic or glass with the REC media. For *in vivo* animal studies, mouse cystic renal tissue was provided by the Maryland PKD Research and Translational Core Center. *Pkd2^fl/fl^ Pax8rtTA TetOCre* mice on a C57BL/6 background, which received doxycycline through intraperitoneal injection of doxycycline suspended in sterile water (50 μg of doxycycline/kg of body weight/day) for two days on P10 and P11. On P15 or P25, mice were euthanized according to IACUC protocols and kidneys were dissected and fixed for processing.

### 3D tubuloid culture

Aliquots of basement membrane construct, growth factor reduced Matrigel (Corning 354230), from −20°C were thawed overnight at 4°C to prepare for tubuloid culture plating. Prechilled 24 well plates and P200 pipet tips were used to set up cultures by first plating 200 μL Matrigel to cover the surface of each well. The bottom layer was allowed to polymerize for 1-2 hours at 37°C. Tubuloid media was made from 37 mL of Basic tubuloid media (490 mL DMEM/F12, 1% Pen/Strep (Sigma P4333), 1% ITS (Gibco 51500-056)) filtered with a 0.22 μm Steriflip, 5 mL fetal bovine serum (Sigma F0926), and a growth factor cocktail of 40 ng/mL recombinant human hepatocyte growth factor (HGF) [Invitrogen PHG0324], 20 ng/mL recombinant human epidermal growth factor (EGF) [Sigma E9644], and 8.8 ng/mL recombinant human basic fibroblast growth factor (FGF) [Sigma FO291]. Mouse glial derived neurotrophic factor (GDNF) (20 ng/mL) [Sigma SRP3200] was added to tubuloid media for pulse media. Following polymerization, cells (*Pkd2* clone #125*, and Pkd1 clones #302, and #313)* from 37°C 2D culture (∼200,000 cells per well), were pelleted using centrifugation, and resuspended in 400 μL of tubuloid media with GDNF and 100 μL of cell suspension was plated on top of first polymerized Matrigel layer. For some limited experiments primary tubule fragments from *Pkd2^fl/fl^, Pax8rtTA, TetOCre, +mTmG* were plated into the 3D culture as described previously^14^. Cells were allowed to attach and settle on Matrigel by incubating at 37°C for 1-2 hours. Finally, the “sandwich” was formed when another 150 μL of Matrigel basement membrane construct was added to cover the primary tubule preparations. The top layer was allowed to polymerize and then cultures were covered with 2 mL tubuloid media with GDNF for 24-36 hours. Tubuloid media (without GDNF) was exchanged every other day by replacing 2 mL of media each time throughout the culture period. Cultures were typically maintained for 14 days. In some experiments we used the ERM phosphorylation inhibitor compound NSC668394 (Sigma 341216) and the Pan PKC inhibitor GO6983 (Med Chem Express HY-13689) which were added directly to the media.

Individual structures were tracked with 4X or 20X brightfield, while population tracking data was acquired with 4X brightfield. Positions for both levels of structure tracking were saved using Metamorph MultiDimensional Acquisition (7.8.12.0) on an Olympus IX83 epifluorescence scope with ASI motorized stage.

### 3D Culture Image Quantification

Images were analyzed in FIJI (ImageJ, enhanced distribution). For TIF images, pixel-to-micron calibration was applied (0.6154 pixels/µm for 4X images acquired on the IX83 Olympus microscope, or as extracted from file metadata). Measurements included area, perimeter, and circularity. Binary images were generated using a custom macro (*Particle Detect.ijm*), which retained only in-focus structures. Fragmented structures were reassembled using binary closing and manual filling tools, while incomplete objects and noise were removed. Completed binary images were saved as TIFFs. Quantification was performed using *Analyze Particles*, with size thresholds adjusted as appropriate. Data outputs were exported to spreadsheets, and overlay functions were used to confirm structure detection. Each image was processed independently, beginning with macro execution.

### Immunostaining

Two dimensional cell cultures were maintain until a few days past confluency to ensure polarization on glass bottomed 12 or 6 wells plates, then cells were fixed with 3% paraformaldehyde (Electron Microscope Solutions 15714-S), permeabilized with 0.1% triton (Sigma T-9284), and blocked in 1% BSA (Sigma A6003) as per standard protocols. Following blocking, polarized cells were incubated overnight at 4°C with primary antibodies in 0.1% BSA in 1X PBS. After washing the primary antibody with PBS, the culture was incubated with secondary antibody in 0.1% BSA. Vectashield Mounting Media with DAPI (Vector Labs H1200) was then added with a cover slip. Samples were imaged on a Nikon W-1 Spinning Disk or the Olympus IX83 inverted imaging system. Primary antibodies used: anti-Ezrin (rabbit, CST 3145), anti-Ezrin (mouse, Sigma 3c12), anti-zonula occludens (ZO) 1 (rat, Santa Cruz sc-33725), anti-NaKATPase (mouse, Millipore 05-369), anti-PKC (rabbit CST 9368) anti-phospho PKC (rabbit CST 2060), and anti-AQP2 (chicken, gift from JB Wade^54^), anti-KI67 (rabbit, CST 9129). Secondary antibodies used: Goat anti-rabbit (Invitrogen A-21429), Goat anti-rat (Invitrogen A-21247), Goat anti-mouse (Invitrogen A-11001), and goat anti-chicken (Invitrogen A21449).

For tissue samples, following dissection, human and mouse kidney samples were fixed in a 3% paraformaldehyde solution, washed in 1X PBS, three times for 10 minutes, and stored in 70% ethanol. Tissue samples were then embedded in paraffin and sectioned. Tissue sections were mounted onto glass coverslips with a gelatin coating solution (50ml distilled water, 0.25g Gelatin, 25mg Chrome Alum) then deparaffinated and placed in a heat induced epitope retrieval (HIER) solution, pH 8.0 (1mM Tris (American Bioanalytical AB02000-01000), 0.5mM EDTA (Sigma E5134) with 0.02% SDS (American Bioanalytical AB01920-00500). Samples were warmed in HIER solution with SDS to 100°C, then transferred to a 100°C water bath for 15 minutes, then washed with distilled water and 1XPBS. Samples were treated with 2-3 drops of Image-iT FX Signal Enhancer (Molecular Probes 136933), then blocked in Incubation Media [1% BSA (Sigma A7638), 0.1% Tween 20 (Bio-Rad 170-6531), 0.02% sodium azide (Sigma S2002),1X PBS (Bio-Rad #161-0780)] with 1% donkey serum (Sigma D9663) for another 15 minutes. Primary antibody (see above) was added in incubation media and serum solution and incubated overnight, then washed four times in 1X PBS. Secondary antibodies (see above) were then added in the dark for two hours and again washed four times in 1X PBS. Following washes, sectioned samples were mounted with Vectashield Mounting Media [Vector Labs H1000], and sealed with nail polish. Samples were imaged on a Olympus IX83 inverted imaging system.

For 3D cell / tubuloid immunostaining we either stained the structures *in situ* or removed them from the Matrigel before processing. For structures in the Matrigel, tubuloids /spheroids were fixed in Matrigel basement membrane by first removing media and washing the culture with prewarmed 1X PBS, three times. Following removal of PBS wash, wells were treated with 1:20 collagenase A (0.1 g/mL; Sigma C2139) and incubated at 37°C for 10 minutes. Enzyme solution was removed and washed again with 1X PBS gently. Tubuloids /Spheroids were fixed in Matrigel with 3% paraformaldehyde (Electron Microscopy Sciences 15714-S) in 1X PBS by rocking plate at room temperature for 30 minutes. Fixative was removed and washed three times with 1X d-PBS. The plate was stored overnight at 4°C in 1X PBS following the fixation steps. Next, it was brought to room temperature and permeabilized by adding 0.025% saponin (Sigma S4521) in 1X PBS, rocking for 30 minutes at room temperature. The 1X PBS washes were repeated and blocked with 1% fish skin gelatin (Sigma G7765) in dH_2_0 for two hours at room temperature. The primary antibodies were added overnight in the fish skin gelatin solution, rocking overnight at 4°C. The following day, the plate was allowed to re-equilibrate to room temperature to reduce the loss of structures during washing. The cultures were washed three times with room temperature 1X PBS. The fluorescent secondary antibody, was added in 1% fish skin gelatin in dH_2_O and incubated while rocking at room temperature for two hours. The culture was washed again for three times with 1X PBS, and the sample was covered with mounting media (Vector Labs H1200). Fixed culture plates were imaged on inverted Olympus IX83 imaging system.

To extract the Tubuloids and Spheroids from the 3D culture system, cultures were rinsed three times with 2 ml cold PBS following media removal, and 1 ml of Corning Cell Recovery Solution (#354253) was added per well. Plates were sealed with parafilm and incubated overnight at 4 °C. Structures were collected using wide-bore BSA-coated pipets into BSA-coated, ice-cold 15 ml conical tubes. Wells were optionally rinsed with an additional 2 ml of recovery solution, which was combined with the initial suspension. Tubes were inverted gently 2–3 times and centrifuged at 1200 rpm for 5 min at 8 °C. Pellets were resuspended in 1 ml cold PBS, centrifuged again under identical conditions, and supernatants reduced to 100 µl (200 µl for two coverslips). Structures were resuspended and plated onto Cell-Tak–coated coverslips (100 µl suspension per coverslip) and allowed to settle for 15 min at room temperature. Excess liquid was carefully removed and retained. Coverslips were washed three times with 2 ml PBS, with each wash collected to recover unattached structures for downstream applications. Coverslips were then fixed in 3% paraformaldehyde (1 ml/well in PBS) for 30 min at room temperature, followed by three PBS washes. Permeabilization was carried out with 0.025% saponin in PBS for 30 min, and coverslips were washed three times with PBS. Blocking was performed with 1% fish skin gelatin in PBS for 2 h at room temperature. Primary antibodies were diluted in blocking buffer, rotated for 30–90 min, and applied to coverslips inverted onto 20–25 µl antibody droplets on parafilm. Samples were incubated overnight at 4 °C in a humidified chamber. The following day, secondary antibodies were diluted in blocking buffer and incubated with samples for 2 h at room temperature in the dark. Coverslips were washed three times with PBS, and excess liquid was removed. For mounting, 10 µl of mounting media was placed on both slide and coverslip; the coverslip was inverted onto the slide to merge the media layers. Edges were sealed with nail polish, and slides were stored at 4 °C (short-term) or –20 °C (long-term).

To quantify Ezrin signal at the membrane in mTmG tubuloids, a background subtracted image was created to remove haze of fluorescence from 3D structures. This was performed in MetaMorph by subtracting (Process) a median filter (Basic Filters) image of the original acquisition for each fluorophore. This resulting images were thresholded and overlayed to create a color combine image of ezrin and either *Pkd2+* or *Pkd2-* tubuloids. Then to quantify the percentage of ezrin signal in each cell of the structures, the composite images were opened in FIJI. A line was drawn around the perimeter of each cell and then the Color Pixel Counter plugin was applied to measure the percentage of overlap color between ezrin and *Pkd2+* (n=32 cells) or *Pkd2-* (n= 40 cells) signal. The statistical comparison between the ezrin signal in *Pkd2+* and *Pkd2-* cells was done by student’s two-tailed t-test.

### Quantitative PCR

cDNA preparation and RTPCR protocols: cDNA was prepared from RNA with the SuperScript III First-Strand Synthesis System for RT-PCR (Invitrogen 18080-051) using the manufacturer’s protocol. Reactions (4 ng cDNA) were run in 96-well plates (Life Technologies 4483485) using manufacturer’s protocol for PowerUp SYBR Green (Life Technologies A25742).

HuEzrin Forward Primer: 5’-CTCGGCGGACGCAAG-3’
HuEzrin Reverse Primer: 5’-CATTGATTGGTTTCGGCATTTTC-3’
HuGAPDH Forward Primer: 5’-GTCTCCTCTGACTTCAACAGCG-3’
HuGAPDH Reverse Primer: 5’-ACCACCCTGTTGCTGTAGCCAA-3’
Mouse Ezrin Forward Primer: 5’-TCCAGTTTAAATTCCGGGCC-3’
Mouse Ezrin Reverse Primer: 5’-TGACTTGCAGGAAGAAGAGC-3’
Mouse GAPDH Forward Primer: 5’-CTTTGGCATTGTGGAAGGGC-3’
Mouse GAPDH Reverse Primer: 5’-TGCAGGGATGATGTTCTGGG-3’

### Western blot protocol

For Cell and Tubuloid western blots, samples were lysed with Deoxycholic Acid RIPA buffer (1% deoxycholic acid, 1% triton X-100, 0.1% SDS, 150mM NaCl, 1mM EDTA, 10mM Tris HCl pH 7.5) with 1:10 protease inhibitor (Sigma P-8340). Western blot of tissue samples of freshly dissected kidney tissue were first homogenized using the BeadBug™6 (Benchtop Scientific) using 3.0 mm beads (Benchmark Scientific, #D1032-30) with the Deoxycholic Acid RIPA lysis buffer. Cell /Tubuloid /tissue lysate was rotated in the cold room at 4°C for 30 minutes and then centrifuged for 15 minutes at 14000 rpm. Supernatant was collected and protein abundance was quantified by bicinchoninic acid (BCA) assay (Thermo Scientific 23225); pellets were saved and stored at −80°C. Samples were heated with 5X laemmeli buffer with sodium dodecyl sodium (SDS) and 10% 2-beta-mercaptoethanol for 30 minutes at 37°C. Samples were then loaded on 10% stain free gels (BioRad 4568033) with kaleidoscope marker (BioRad 161-0375) and run for 50 minutes at 200 V. Before transfer, gels were crosslinked using UV (BioRad ChemiDoc MP Imaging System) and imaged to quantify total loaded protein (LC) for normalization of protein abundance. Gels were then transferred onto 0.2% nitrocellulose membranes (BioRad 1704158) using semidry BioRad Trans-blot Turbo System. Membranes were blocked in 5% Milk in 1X TBS-T and primary antibodies were incubated overnight in 2.5% milk in 1X TBS-T at 4°C. Blots were washed three times in 1X TBS-T, then incubated with secondary antibody (goat anti-rabbit HRP or goat anti-mouse HRP; Jackson Immuno-Research Laboratories 111035144) in 2.5% milk in 1X TBS-T for 1 hour, rocking, at room temperature. Blots were washed again three times with 1X TBS-T and developed in Super Signal ECL (Thermo Scientific 34577). Blots were developed using Biorad Chemidoc Imaging machine and quantified using Image Lab (BioRad Version 6.0.1 build 34). Normalization was done by calculating the total protein loaded in each lane using the BioRad Stain-Free Gel System. Statistical comparisons of density measurements from western blots were done with the Student’s t-test for pair-wise comparisons (Prism 8, GraphPad, USA). All reported means are ± standard error of the mean (SEM).

### Protein interactions studies

Antibodies used for the immunoprecipitation experiments: anti-flag M2 magnetic beads (Sigma M8823), anti-HA beads (mouse, Sigma, A2095), anti-Ezrin (rabbit, CST 3145), anti-Myc (rabbit CST 2276), anti-HA (mouse, Sigma H9658), anti-Polycystin 1 -E8, and anti-Polycystin 1 #45 (both rabbit, F. Qian / PKD-RRC).

#### Exvivo Immunoprecipitation with endogenous proteins

Immunoprecipitation (IP) studies of HA-tagged PC1 and PC1V were performed on mouse kidney epithelial cells isolated from P4-P5 pups and cultured (as described above) for 4 days. Samples were homogenized in lysis buffer (25 mM Na phosphate [pH 7.2], 150 mM NaCl, 1 mm EDTA, 10% glycerol, 1% Triton X-and protease inhibitor cocktail [Roche]) for 1h at 4°C. Approximately 2mg of clear protein lysates were used for IP with anti-HA-agarose beads. The IP reactions were loaded on 4-12% Bis-tris or 3-8% Tris-acetate-SDS-polyacrylamide precast gels (Invitrogen) and transferred to PVDF membranes (Bio-Rad). The membranes were probed overnight at 4°C with primary antibodies diluted in TBST containing 5% non-fat dry milk or 3% BSA. Protein detection was achieved by chemiluminescence using HRP-conjugated secondary antibodies and ECL reagent (Amersham).

#### Immunoprecipitation with transfected constructs

Constructs: FL-PC1 with C-terminal flag (AF20), FL-PC1V with C-terminal flag, FL-PC2 with N-terminal Myc (OF2-3) and CT-PC2 with N-terminal Myc (only final TM and C-terminal tail, HBF292, all were generous gifts from F. Qian^13^). The Ezrin construct was created by cloning Ezrin (EE Dixon) using the Ezrin GBlock inserted into a TOPO vector using the Zero Blunt TOPO PCR Cloning Kit (ThermoFisher K280020). The Ezrin insert was then cut from the TOPO vector and ligated into pCS2+ using T4 DNA ligase (NEB M0202). Constructs were transfected into HEK293T cells (media: 1X DMEM (Gibco 11965-092) with 10% FBS (Sigma F0926), 1% PS (Gibco 15140-122), 1% Glutamax (Gibco 35050-061)) in a 6 well plate using Xtreme Gene HP (Roche 06 366 236 001) according to manufacturer’s protocol. Following 36 hours, transfected cells are lysed using deoxycholic RIPA lysis buffer (described above) with 1:100 protease inhibitor (Sigma P8340) and then quantified using BCA assay via manufacturer’s protocol. Antibody conjugated Magnetic beads were combined with: (a) deoxycholic RIPA buffer, for PC2 immunoprecipitations (up to 1 mL) or (b) Polycystin Lysis / wash buffer (PLB) for PC1 immunoprecipitations, plus cell lysate (50-100 μg protein) or IgG (Mouse Santa Cruz sc-2025) overnight and rotated at 4°C. The next day, a magnet was used to collect beads and wash with HEENG buffer pH 7.4 for PC2 immunoprecipitations (20mM HEPES pH 7.6, 125mM NaCl, 1mM EDTA, 1mM EGTA, 100 mL glycerol) or PLB PC1 immunoprecipitations (50 mM Tris-HCL pH 7.5, 150 mM NaCl, 1% NP-40) four times for 5-10 minutes each. Following the last buffer wash, beads were again collected using the magnet and elution buffer (1:1 2X laemelli buffer with 10% beta-mercaptoenthanol to deoxycholic RIPA buffer or PLB) was added and samples were eluted at 42°C for 45 minutes, disrupting the sample every 15 minutes. Using the Western blot protocol described above, eluted samples were run with 1% input on 4-20% gels (BioRad 456-1094).

## Acknowledgments

The authors gratefully acknowledge the Maryland PKD Research and Clinical Core Center (as part of the PKD RRC) for providing valuable reagents and resources. The work of O.M.W. was supported by NIDDK grants P30DK090868, U54DK126114, and a grant from the PKD foundation (Research Grant 229G18a). EED was supported by F31DK117579, AZ was supported by the T32DK098107, and VHK was supported by F32DK124960. The views expressed in this manuscript are those of the authors and do not necessarily represent the views of the National Institutes of Health or any other funding source.

## Author contributions

Eryn E. Dixon: concept / design; acquisition; analysis; interpretation of data; substantively revised manuscript

Ava Zapf: concept / design; acquisition; analysis; interpretation of data

Victoria Halperin Kuhns: acquisition; analysis; interpretation of data

Denis Basquin: acquisition; analysis

Rachel Park: acquisition

Richard Coleman: acquisition

Lidiya Franklin: acquisition; analysis

Allison C. Lane-Harris: acquisition

Alexis Hofherr: concept / design

Michael Kottgen: concept / design

Feng Qian: analysis; interpretation of data

Paul A. Welling: concept / design; analysis; interpretation of data; substantively revised manuscript

Terry J. Watnick: concept / design; analysis; interpretation of data; substantively revised manuscript

Owen M. Woodward: concept / design; acquisition; analysis; interpretation of data; drafted manuscript; substantively revised manuscript

