## Supplemental Figures for "Extra-ciliary role for polycystins in regulation of Ezrin and renal tubular morphology"

**A.**

#### Supplemental Figure 1

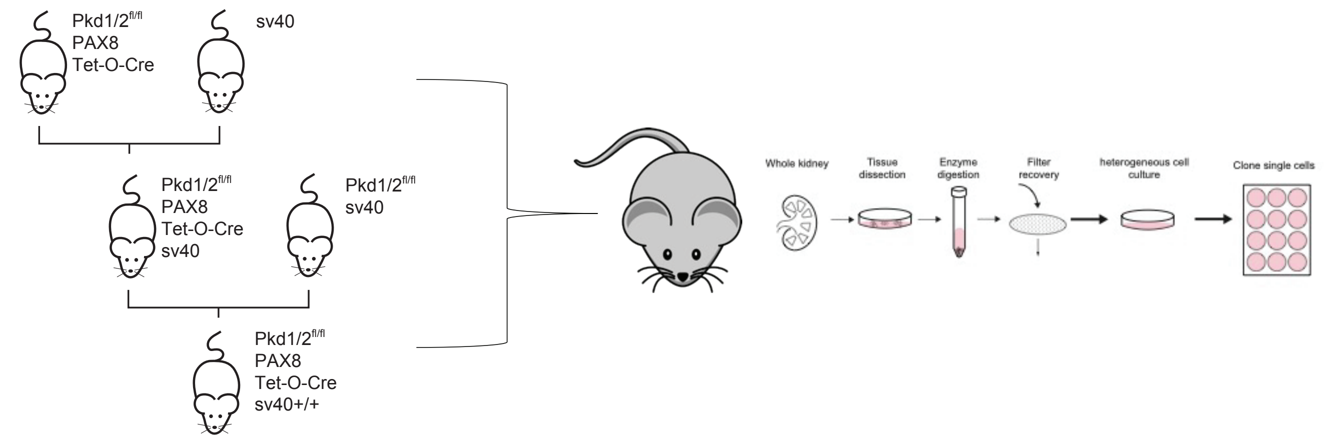


| Clone # | Genotype | origin | sex | Validated KO | 3D validated | Published | Column1 |
|---|---|---|---|---|---|---|---|
| 125 | Pkd2 fl/fl; Pax8+ Cre+; SV40 | medulla | male | X | X | Jung et al. 2023 | |
| 302 | Pkd1 fl/fl; Pax8+ Cre+; SV40 | cortex | male | X | X | | |
| 313 | Pkd1 fl/fl; Pax8+ Cre+; SV40 | medulla | male | X | X | | |

(A) Work flow for the creation of the immortalized mouse renal epithelial cells with inducible knock-out of Pkd1 or Pkd2 (see methods for additional details)

**B.**

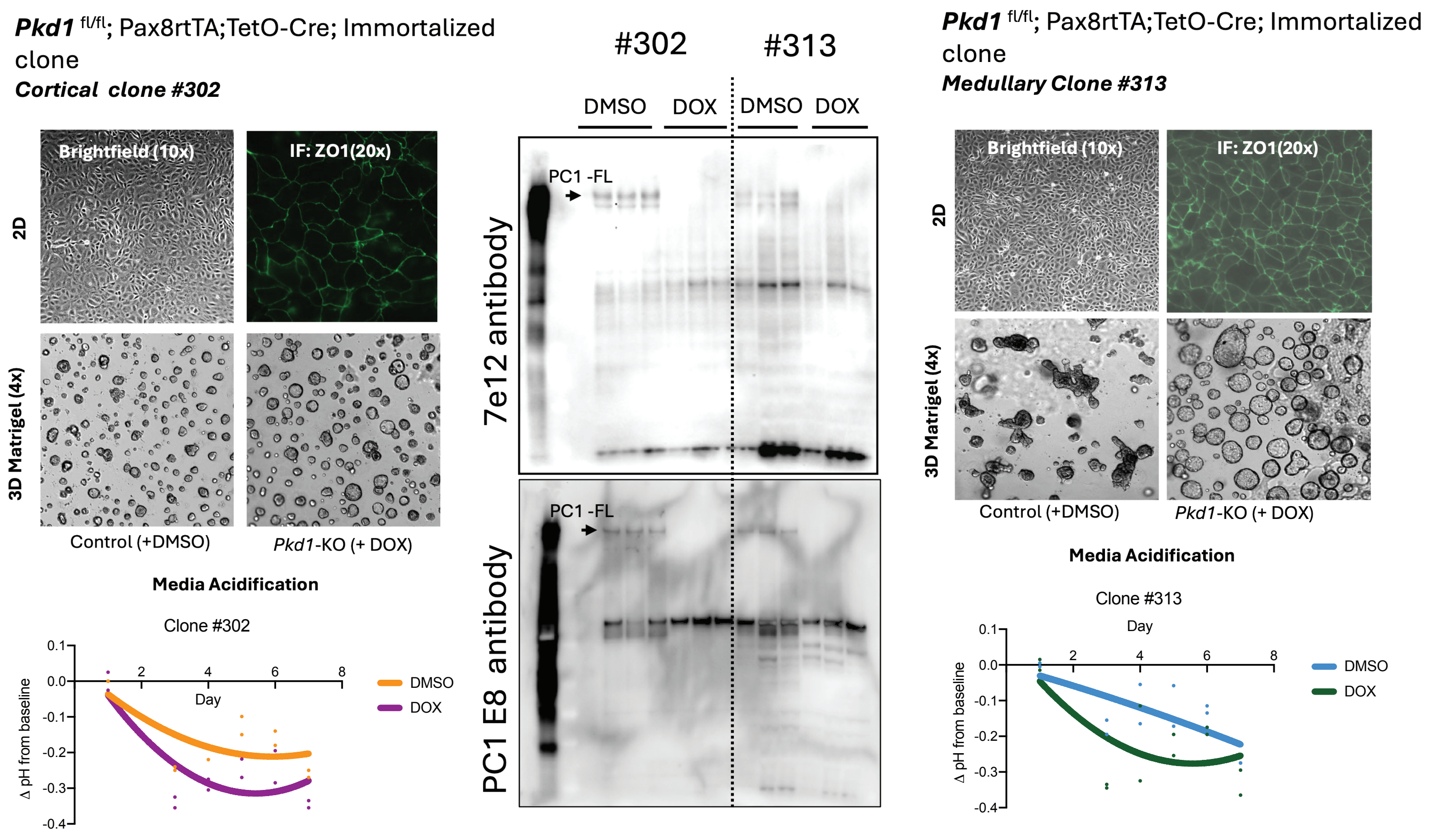


(B) Characteristics and function data for clone #302 and #313: 2D and 3D micrographs, IF of junctional marker ZO1, and the tracking of media pH of time. Media acidification is a hall mark of cells with Pkd1 KO. See methods for additional details

A

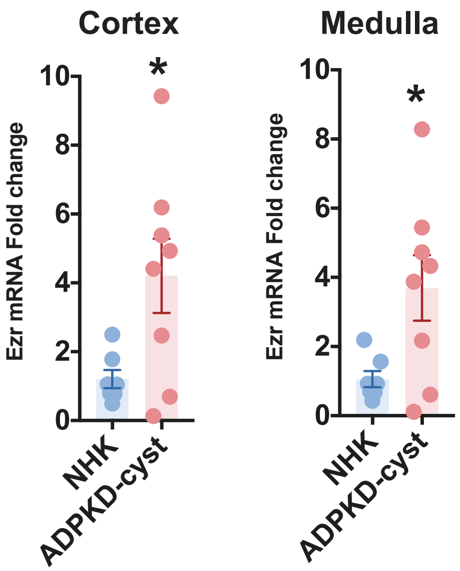


#### EZR gene expression in adult human normal and ADPKD ESRD kidneys

(A) Real Time rt-PCR of samples from normal human kidney tissue (NHK, cortex/medulla) and cyst tissue (cortex-/medulla) dissected from ADPKD kidneys (n=7/8; +/- SE; Student T test, two tailed, p<0.05).

B

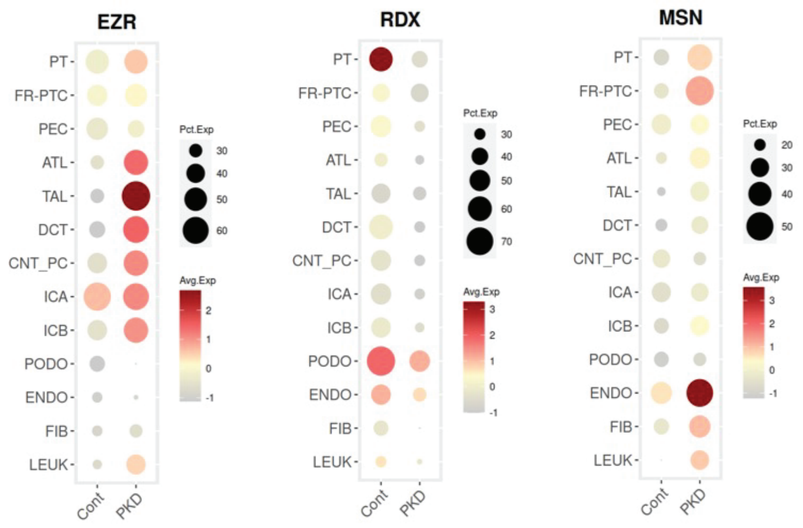


#### ERM gene expression in adult human normal and ADPKD ESRD across the nephron.

(B) Gene expression across different cell types of the human kidney for Ezr (Ezrin), RDX (Radixin), and MSN (Moesin). Data from :

Muto, Y., Dixon, E.E., Yoshimura, Y. et al. Defining cellular complexity in human autosomal dominant polycystic kidney disease by multimodal single cell analysis. Nat Commun 13, 6497 (2022). https://-doi.org/10.1038/s41467-022-34255-z

C

| Gene_symbol | Annotation | PTS1 | PTS2 | PTS3 | DTL1 | DTL2 | DTL3 | ATL | MTAL | CTAL | DCT | CNT | CCD | OMCD | IMCD |
|---|---|---|---|---|---|---|---|---|---|---|---|---|---|---|---|
| Ezr | ezrin | 252.9 | 223.2 | 117.2 | 309.6 | 302.5 | 418.9 | 283.5 | 303.1 | 378.2 | 361.4 | 469.8 | 449.9 | 293.2 | 235.6 |
| Rdx | radixin | 60.4 | 64.5 | 42.5 | 117.6 | 118.6 | 103.1 | 219.5 | 76.3 | 90.2 | 67.9 | 72.1 | 67.3 | 84.8 | 109.6 |
| Msn | moesin | 63.1 | 46.4 | 45.9 | 333.4 | 308 | 200.2 | 298.8 | 67.7 | 111 | 40.5 | 26.4 | 14.8 | 26.4 | 3.5 |

#### ERM gene expression in adult mouse kidneys.

(C) Gene expression across different renal tubule segments of themouse kidney for Ezr (Ezrin), RDX (Radixin), and MSN (Moesin). Data from :

Chen L, Chou CL, Knepper MA. A Comprehensive Map of mRNAs and Their Isoforms across All 14 Renal Tubule Segments of Mouse. J Am Soc Nephrol. 2021 Apr;32(4):897-912. doi: 10.1681/ASN.2020101406.

Supplemental Figure 3

**A**

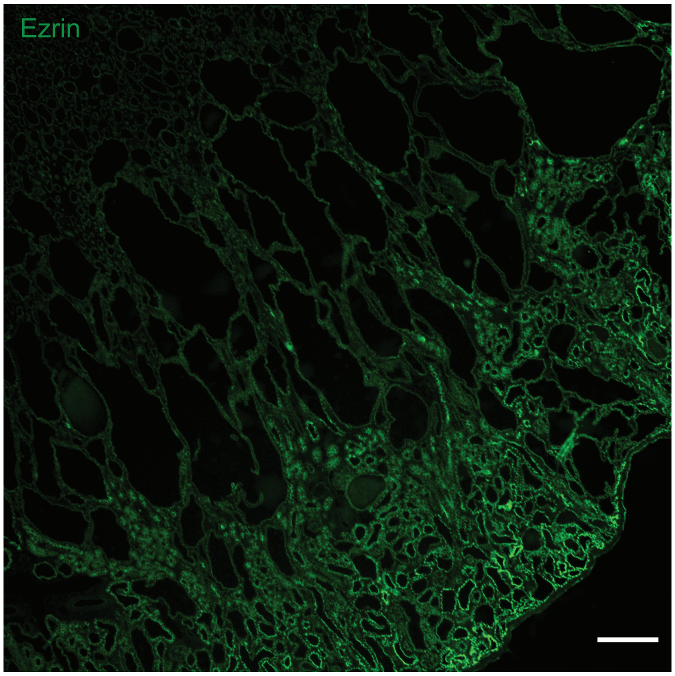


Ezrin in *Pkd2* inactivation in vivo model (P25) mice

(A) Ezrin (green) immunofluorescence in Pkd2fl/fl Pax8 rtTA TetOCre male mice collected fifteen days (P25) following doxycycline. While ezrin does not seem to localize to the apical membrane of medullary cysts, there is appropriate signal in noncystic proximal tubules. Magnfication 4X, Scale bar 250 µm.

### Supplemental Figure 4

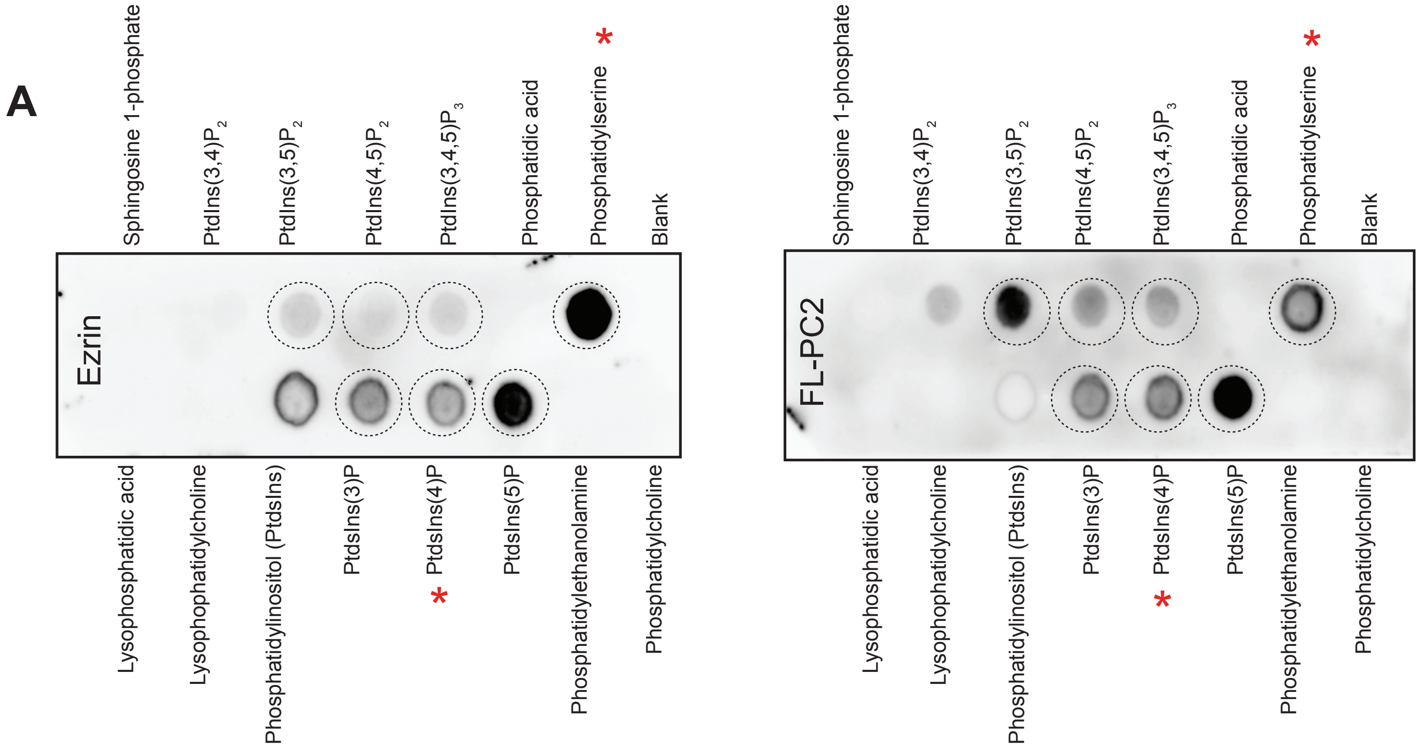


(A) Lipid overlay reveals a similar profile of interacting phosphoinositides (dotted black circles) between over expressed Ezrin and FL-PC2 in HEK293T cell lysate. Lipid overlay assays: PIP strips (Invitrogen P23751) were blocked in blocking solution (1X TBS-T with 3% fatty acid free BSA (Sigma A7030)) for one hour at room temperature. Following blocking, 50 µg/mL of protein lysate (Ezrin, PC2FL, PC2CT, or untransfected control) in blocking solution was added to the strip membrane overnight to rock at 4$^{O}$C. The next day, PIP strips were rocked for an additional hour at room temperature and then washed three times with 1X TBS-T, followed by blocking in 5% milk for one hour. Primary antibodies (1:1000 Ezrin or 1:1000 Myc) were added to the PIP strips and incubated for two hours at room temperature in 2.5% milk in 1X TBS-T. Then the PIP strips were incubated with secondary antibodies (goat anti rabbit or mouse HRP 1:5000) and developed with chemiluminescence.

(*) Phosphoinositides previously were shown to interact with PC1 PLAT domain (Xu Y, Streets AJ, Hounslow AM, Tran U, Jean-Alphonse F, Needham AJ, Vilardaga JP, Wessely O, Williamson MP, Ong AC. The Polycystin-1, Lipoxygenase, and α-Toxin Domain Regulates Polycystin-1 Trafficking. J Am Soc Nephrol. 2016 Apr;27(4):1159-73. doi: 10.1681/ASN.2014111074. Epub 2015 Aug 26.).
